# Genome sequence and cell biological toolbox of the highly regenerative, coenocytic green feather alga *Bryopsis*

**DOI:** 10.1101/2023.11.22.568388

**Authors:** Kanta K. Ochiai, Daiki Hanawa, Harumi A. Ogawa, Hiroyuki Tanaka, Kazuma Uesaka, Tomoya Edzuka, Maki Shirae-Kurabayashi, Atsushi Toyoda, Takehiko Itoh, Gohta Goshima

## Abstract

Green feather algae (Bryopsidales) undergo a unique life cycle in which a single cell repeatedly executes nuclear division without cytokinesis, resulting in the development of a thallus (> 100 mm) with characteristic morphology called coenocyte. *Bryopsis* is a representative coenocytic alga that has exceptionally high regeneration ability: extruded cytoplasm aggregates rapidly in seawater, leading to the formation of protoplasts. However, the genetic basis of the unique cell biology of *Bryopsis* remains poorly understood. Here, we present a high-quality assembly and annotation of the nuclear genome of *Bryopsis* sp. (90.7 Mbp, 27 contigs, N50 = 6.7 Mbp, 14,034 protein-coding genes). Comparative genomic analyses indicate that the genes encoding BPL-1/Bryohealin, the aggregation-promoting lectin, are heavily duplicated in *Bryopsis*, whereas homologous genes are absent in other Ulvophycean algae, suggesting the basis of regeneration capability of *Bryopsis*. *Bryopsis* sp. possesses >30 kinesins but only a single myosin, which differs from other green algae that have multiple types of myosin genes. Consistent with this biased motor toolkit, we observed that the bidirectional motility of chloroplasts in the cytoplasm was dependent on microtubules but not actin in *Bryopsis* sp. Unexpectedly, most genes required for cytokinesis in plants are present in *Bryopsis*, including those in the SNARE or kinesin superfamily. Nevertheless, a kinesin crucial for cytokinesis initiation in plants (NACK/Kinesin-7II) is hardly expressed in the coenocytic part of the thallus, possibly underlying the lack of cytokinesis in this portion. The present genome sequence lays the foundation for experimental biology in coenocytic macroalgae.

**Significance statement:** The exceptionally coenocytic body and remarkable regeneration ability of *Bryopsis* have attracted biologists for years. However, molecular biological tools remain underdeveloped, partly due to the lack of genome information. Here, we report high-quality assembly and annotation of the genome, providing a crucial resource for experimental biology and genomics studies of *Bryopsis*. Furthermore, comparative genomic analysis reveals a unique gene repertoire that possibly underlies the highly regenerative coenocytic body.

## Introduction

Eukaryotic cells are typically characterised by a single nucleus at the centre of the cytoplasm. However, some exceptions exist. For example, red blood cells are anucleated. Multinucleated cells have also been observed in a variety of species. In animals, the syncytium in *Drosophila* embryos and muscle cells in mammals have been extensively studied in cell and developmental biology, for example for the mechanisms of nuclear positioning and synchronised nuclear division (Kwon and Scholey, 2004; Padilla et al., 2022). In flowering plants, seed endosperm undergoes repeated mitotic nuclear divisions without cytokinesis after double fertilisation, forming a large multinucleated cell called ‘coenocyte’ (Ali et al., 2023). Many species of marine macroalgae (seaweeds) possess multinucleated cells in their body (Graham et al., 2008). An extreme situation is seen in green feather algae; the thalli of *Caulerpa* or *Bryopsis* develop and reach over 10 cm in length with characteristic side branches, but strikingly, there are no cell walls to separate the numerous nuclei (Mine et al., 2008). This coenocytic feature raises many evolutionary and cellular biology questions, such as how the characteristic features evolve specifically in this algal lineage or how intracellular components are organised in the extremely large cytoplasm (Mine et al., 2008; Umen and Herron, 2021). Non-uniform distribution of transcripts might partly contribute to cytoplasmic organisation in coenocytes (Ranjan et al., 2015). However, the underlying mechanism remains poorly understood, partly because of the lack of an experimental model system in which genetic and molecular biological tools can be instantly applied. As the first step, it is critical to understand the genome sequences and gene repertoires of these species.

Among green feather algae, *Bryopsis* has garnered special attention for its remarkable regenerative capabilities in laboratory settings: cytoplasm extruded from mature thalli is rapidly clustered and transformed into protoplasts, followed by thallus development under the laboratory culture condition (Ikeuchi et al., 2016; Kim et al., 2001; Pak et al., 1991; Tatewaki and Nagata, 1970). This amazing regeneration ability, undergoing ‘life without a cell membrane’ (Kim et al., 2001), might be critical for this single-celled organism when they are physically damaged, for example by predators (Zan et al., 2019). Regarding the factors required for regeneration, Kim and colleagues found that the aggregation of the extruded cytoplasm is facilitated by the F-type domain-containing lectin termed Bryohealin (also called BPL-1) in *B. plumosa* (Kim et al., 2006). The BPL-1-like protein similarly facilitates aggregation in *Bryopsis hypnoides* (Niu et al., 2009). Aggregation is inhibited by N-acetyl-D-glucosamine and N-acetyl-D-galactosamine, which possess high affinity to BPL-1 (Kim et al., 2006; Niu et al., 2009; Yoon et al., 2008). Three other types of lectins, BPL-2 (Han et al., 2010a), BPL-3 (Han et al., 2010b), and BPL-4 (Han et al., 2012), have been also identified in *Bryopsis*, which bind to the above two sugars (BPL-3/4) or D-mannose (BPL-2). Since extremely high regeneration ability is a unique feature of *Bryopsis*, an interesting scenario would be that some of these lectins uniquely evolved in *Bryopsis*.

The taxon Chlorophyta, to which most green algae belong, exhibits remarkably varied body plans (Del Cortona et al., 2020; Gulbrandsen et al., 2021; Hou et al., 2022; Leebens-Mack et al., 2019). Green microalgae, such as the model species *Chlamydomonas reinhardtii*, are generally unicellular with a single nucleus, whereas Dasycladales, including the classical cell biology model organism *Acetabularia*, is unicellular with a single nucleus but with extremely large cytoplasm (up to 10 cm). Ulvales species have canonical multicellular bodies that are made of mono-nucleated cells separated by cell walls, while Cladophorales is multicellular with multiple nuclei per cell. Several genome sequences of green algae are available, including those of the coenocytes *Caulerpa lentillifera* and *Ostreobium quekettii* (Arimoto et al., 2019; Hanschen and Starkenburg, 2020; Iha et al., 2021). However, genomic information is lacking for the family Bryopsidaceae, which includes the genus *Bryopsis*. Moreover, the gene repertoire that possibly characterises coenocytic cells has not been extensively investigated yet. In this study, we present the first and high-quality genome sequences of *Bryopsis* species (registered as *Bryopsis* sp. KO-2023). We then report the cell biological toolbox of *Bryopsis* and other green algae.

## Results and Discussion

### Characterisation of *Bryopsis* species isolated on Sugashima Island, Japan

We isolated two *Bryopsis*-like specimens from an outdoor tank at Sugashima Marine Biological Laboratory (Fig. 1A). Sequencing of the rDNA ITS locus showed > 99.5% identity in 437 base pairs (bp) with that of a *Bryopsis* species registered in the database (line name: HIRO:HIRO-MY 77087). DNA staining showed that multiple nuclei were distributed in the cytoplasm of the main axis, confirming the coenocytic feature (Fig. 1A, middle). High regeneration ability was also confirmed. When the cytoplasm was squeezed out, the extrusion quickly aggregated and transformed into a membrane-encircling protoplast, followed by tip growth (Fig. 1B, Movie 1). Furthermore, this process was suppressed by N-acetyl-D-glucosamine (Fig. S1A) (Kim et al., 2006; Niu et al., 2009).

**Figure 1.**
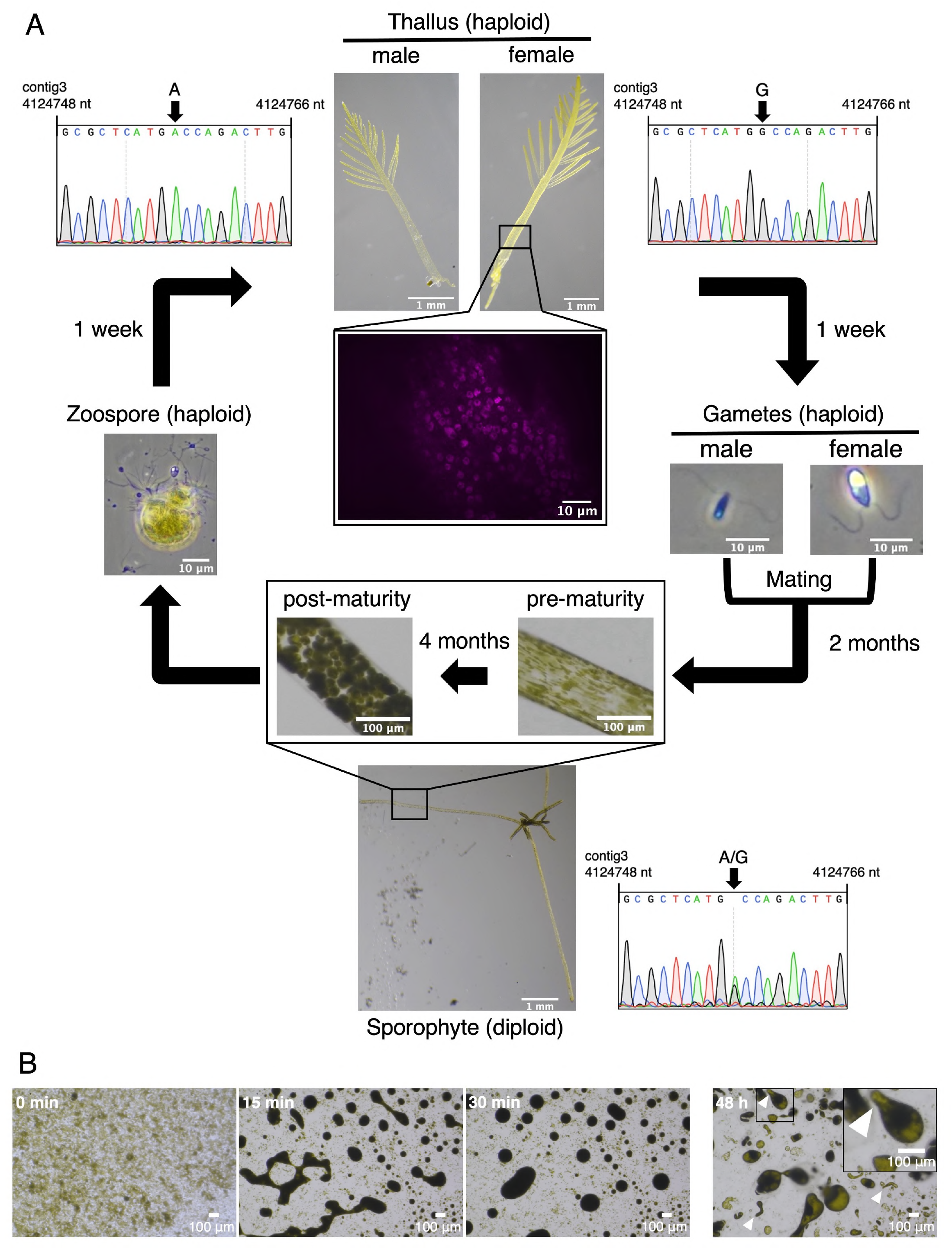
Life cycle and regeneration of *Bryopsis* collected on Sugashima Island. (A) Life cycle of *Bryopsis*. Images are derived from *Bryopsis* sp. analysed in this study. Sequencing indicates a SNP in male and female lines (contig 3: nt 4124748–4124766). Note that both A and G were detected in the sporophyte (diploid). DAPI-stained (magenta) nuclei are shown in the middle. (B) Regeneration of *Bryopsis* sp. after extrusion of the cytoplasm into autoclaved seawater. See also Movie 1. Arrowheads indicate polarised tip growth of regenerated cells.

Next, we tested whether the obtained lines underwent a previously reported life cycle (Tatewaki, 1973). The morphology of the gametes suggested that one line was male and the other was female. Under conditions similar to those used in previous studies, we successfully observed gamete production from both lines, mating of the gametes to generate a sporophyte (diploid), and zoospore generation (Fig. 1A).

We also observed the microtubules and actin filaments using confocal microscopy after immunostaining. They were observed only near the thallus surface, that is, in the cortical cytoplasm, and ran along the main axis of the thallus (Fig. S1B, C). They overlapped largely, but not entirely. The microtubules were not visible after treatment with oryzalin, a microtubule-destabilising drug widely used in land plants. Colocalised actin filaments were also diminished, whereas other short actin bundles remained (Fig. S1D). In contrast, the commonly used actin inhibitor, latrunculin A, completely destroyed actin filaments, whereas microtubule bundles remained intact (Fig. S1E). These observations are largely consistent with those of previous studies using different drugs and epifluorescence microscopy (Menzel and Schliwa, 1986a; Menzel and Schliwa, 1986b).

Based on these observations, we concluded that the collected lines were male and female *Bryopsis*.

### Genome sequences and annotation – nucleus

We extracted RNA and DNA separately from haploid thalli (female) and performed sequencing. A draft nuclear genome was assembled based on the short and long reads. The genome comprised 27 contigs (90.7 Mbp, N50 length 6.7 Mbp) (Table 1). The average coverage was 45× (short reads) and 322× (long reads). The GC content was 45.9%, similar to that of *O. quekettii* (52.4%) and *C. lentillifera* (40.4%) (Table S1).

**Table 1.**
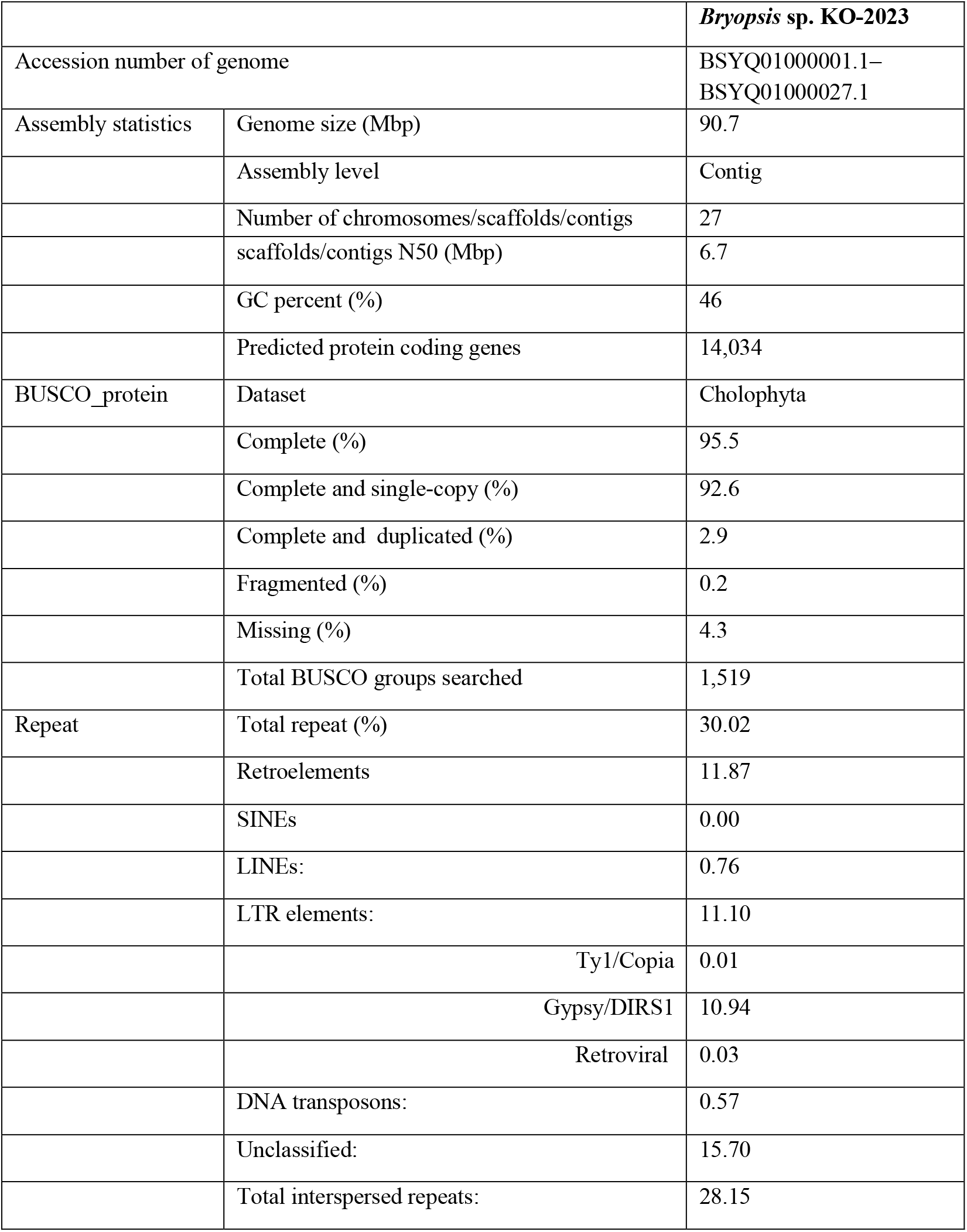
Information of the nuclear genome of *Bryopsis* sp. KO-2023.

|  |  | *Bryopsis* sp. KO-2023 |
| --- | --- | --- |
| Accession number of genome |  | BSYQ01000001.1–<br>BSYQ01000027.1 |
| Assembly statistics | Genome size (Mbp) | 90.7 |
|  | Assembly level | Contig |
|  | Number of chromosomes/scaffolds/contigs | 27 |
|  | scaffolds/contigs N50 (Mbp) | 6.7 |
|  | GC percent (%) | 46 |
|  | Predicted protein coding genes | 14,034 |
| BUSCO_protein | Dataset | Cholophyta |
|  | Complete (%) | 95.5 |
|  | Complete and single-copy (%) | 92.6 |
|  | Complete and duplicated (%) | 2.9 |
|  | Fragmented (%) | 0.2 |
|  | Missing (%) | 4.3 |
|  | Total BUSCO groups searched | 1,519 |
| Repeat | Total repeat (%) | 30.02 |
|  | Retroelements | 11.87 |
|  | SINEs | 0.00 |
|  | LINEs: | 0.76 |
|  | LTR elements: | 11.10 |
|  | Ty1/Copia | 0.01 |
|  | Gypsy/DIRS1 | 10.94 |
|  | Retroviral | 0.03 |
|  | DNA transposons: | 0.57 |
|  | Unclassified: | 15.70 |
|  | Total interspersed repeats: | 28.15 |

Several contigs had a common repeat sequence (CCCTAAA) at the end (Fig. 2A, red bars at the end of contigs). This sequence was identical to the telomeric repeat sequences of *Arabidopsis thaliana* (Richards and Ausubel, 1988), suggesting that they represent the chromosomal end. This repeat was identified at both ends of the five contigs, suggesting that complete sequences of the five chromosomes were obtained in our analysis. In the other eight contigs, the repeat was observed at one end. Provided that this repeat indeed represents telomeric sequences, *Bryopsis* sp. haploid would possess nine or more chromosomes.

**Figure 2.**
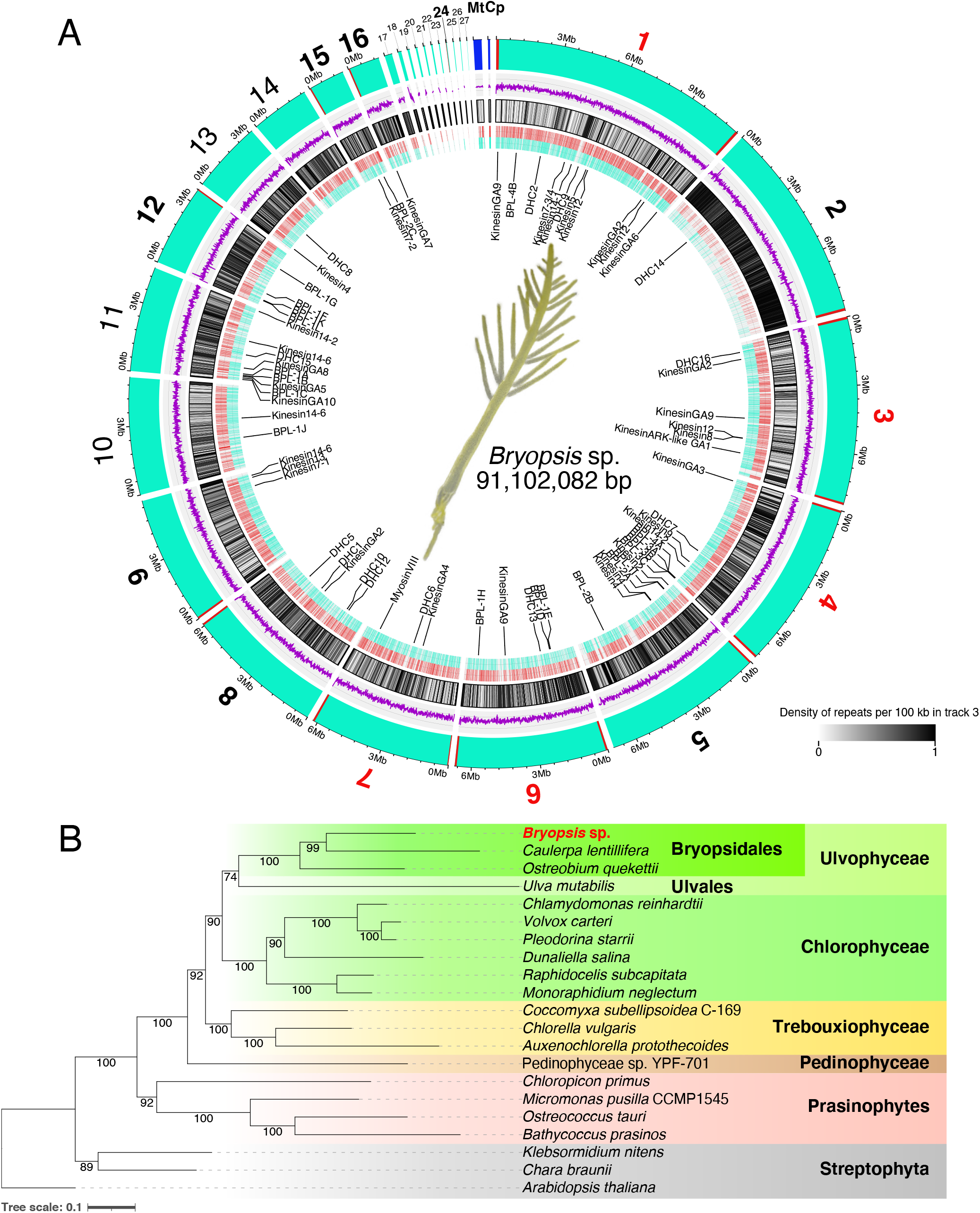
Nuclear and organelle genome assembly. (A) Circos plot of the 27 contigs and organelles assembled from *Bryopsis* sp. (From outmost to innermost lanes) (1) Contigs (cyan) and putative telomeric repeats (red bar, CCCTAAA) are shown. When the repeat was identified in both ends of the contig, the contig number was indicated in red. When just one end had the repeat, the contig was highlighted with a black bold letter. Blue bars indicate organelles of circular genome (mitochondrion: Mt, chloroplast: Cp). (2) Purple lines indicate G/C content per 10,000 bp. Two grey lines indicate 25% and 75%. (3) Black bars present non-telomeric repeat sequences. (4) Red and blue bars indicate genes from Watson and Crick strands, respectively. (5) Genes analysed in this study. (B) Phylogenetic tree of green algal species subjected to KEGG analysis in this study. Maximum Likelihood (ML) tree was constructed with LG+F+R4 selected as the best-fit model and the branch support was estimated with 1,000 ultrafast bootstrap. The bar indicates 0.1 amino acid substitutions per site.

A total of 14,034 protein-coding genes were predicted in 27 contigs (Table 1). BUSCO analysis (protein mode) using the chlorophyta lineage dataset indicated that 92.6% of the single-copy orthologues were recovered, which was higher than those of *O. quekettii* (55.0%) and *C. lentillifera* (67.0%) (Table S1).

These analyses suggest that the nuclear genome of *Bryopsis* sp. was assembled and annotated with high quality compared to many other algal genomes (Hanschen and Starkenburg, 2020).

### Genome sequences and annotation – chloroplast and mitochondrion

The chloroplast genome was assembled into a single circular sequence. The number and identity of protein-coding genes, rRNA, and tRNA, as well as the overall genome size were comparable to those of the reported sequences derived from *B. plumosa* and *B. hypnoides* (Leliaert and Lopez-Bautista, 2015; Lu et al., 2011) (Table 2). Detailed information on the genome, including unique features identified in our line, is provided in the **Supplementary Document**.

**Table 2.** Comparison of the chloroplast and mitochondrial genome.

|  | *Bryopsis*<br>sp. | *Caulerpa*<br>*lentillifera* | *Ostreobium*<br>*quekettii* | *Ulva* sp. | *Chlamydomonas*<br>*reinhardtii* | *Physcomitrium*<br>*patens* | *Arabidopsis*<br>*thaliana* |
| --- | --- | --- | --- | --- | --- | --- | --- |
| Genome | Chloroplast | Chloroplast | Chloroplast | Chloroplast | Chloroplast | Chloroplast | Chloroplast |
| Accession number of genome | LC76890 1.1 | NC_039377.1 | NC_030629.1 | KP72061 6.1 | NC_005353.1 | NC_005087.2 | NC_000932.1 |
| Genome size (Kbp) | 91.7 | 119.4 | 82.0 | 100.0 | 203.8 | 122.8 | 154.4 |
| GC percent (%) | 30.4 | 32.6 | 31.9 | 25.3 | 34.5 | 28.5 | 36.3 |
| Predicted protein coding genes* | 83 | 91 | 78 | 79 | 65 | 85 | 79 |
| rRNA genes* | 3 | 3 | 3 | 3 | 5 | 3 | 4 |
| tRNA genes* | 26 | 28 | 31 | 28 | 29 | 32 | 30 |
| Coding DNA (%) ** | 85.4 | 86.0 | 84.0 | 81.8 | 49.9 | 72.3 | 72.0 |
| Large inverted repeat (>5 kb) | absent | absent | absent | absent | present | present | present |
| Genome | Mitochondrion | Mitochondrion | Mitochondrion | Mitochondrion | Mitochondrion | Mitochondrion | Mitochondrion |
| Accession number of genome | LC76890 2.1 | KX76157 7.1 | NC_045361.1 | KP72061 7.1 | NC_001638.1 | NC_007945.1 | NC_037304.1 |
| Genome size (Kbp) | 356.2 | 209 | 241.7 | 73.5 | 15.8 | 105.3 | 367.8 |
| GC percent (%) | 54.4 | 50.9 | 48.3 | 32.4 | 45.2 | 40.6 | 44.8 |
| Predicted protein coding genes* | 54 | 76 | 54 | 50 | 8 | 42 | 122 |
| rRNA genes* | 3 | 3 | 3 | 2 | 14 | 3 | 3 |
| tRNA genes* | 17 | 20 | 28 | 25 | 3 | 24 | 22 |
| Intron number | 72 | 29 | 47 | 10 | 0 | 26 | 18 |
| Intronic DNA (%) ** | 54.1 | 43.4 | 39.3 | 21.7 | 0 | 28.4 | 8.14 |

\* Duplicate genes were counted as single genes.
\*\* Total gene length, which includes introns, was divided by the entire genome length.

The mitochondrial genome was assembled into a single circular sequence (Table 2). Our sequence substantially diverged from the reported ‘*Bryopsis plumosa*’ sequence (Han et al., 2020). However, our own analysis of the reported sequences indicated that the specimen belonged to the order Ulvales, and not Bryopsidales (Fig. S2). We think that ours represent the first full mitochondrial DNA sequences of *Bryopsis*. The detailed description the genome feature is provided in the **Supplementary Document**.

### Overview of the *Bryopsis* sp. nuclear genome

The availability of high-quality genome allowed us to conduct a high-level comparative genomic study of *Bryopsis*. As comparison, we selected two land plant species and 20 green algal species (5 macroalgae and 15 microalgae), which covered several classes in Chlorophyta (Fig. 2B, Table S2). The genomes of most species have been annotated in high quality, except for *O. quekettii* (Bryopsidales), whose BUSCO value (genome mode) is less than 70% (Table S1).

First, the comparison of the sequences of 10 single copy genes indicated that *Bryopsis* sp. was indeed phylogenetically classified into the order Bryopsidales and was closer to *C. lentillifera* than *O. quekettii* (Fig. 2B) (Del Cortona et al., 2020; Gulbrandsen et al., 2021; Hou et al., 2022; Leebens-Mack et al., 2019). Second, the repeat sequences were surveyed, as they would reflect the phylogeny (Dodsworth et al., 2014). In all three Bryopsidales species, Ty1/Copia-type long terminal repeat (LTR) retrotransposons were scarcely detected (<0.01%), in contrast to their prevalence in *Ulva mutabilis*, *C. reinhardtii*, and land plant (Table S1). The LINEs were also infrequently detected in Bryopsidales. These results are consistent with the phylogenetic tree derived from gene sequences. Third, we provided functional annotation based on KEGG (Kyoto Encyclopedia of Genes and Genomes) and investigated which unigenes are over-or under-represented in *Bryopsis* (Table S3). *Bryopsis* sp. had >10% more unigenes than the average numbers of green algae in several categories, including signal transduction, transport and catabolism, and cell motility (Table S3). This analysis, however, could not be applied to other Bryopsidales species, as their relatively poor gene annotation would result in underestimation of the unigene numbers. We next analysed total numbers of the genes in each category, which would be less sensitive to genome quality. This analysis showed that the genes in the signalling pathway including SnRK2 kinase were expanded in Bryopsidales (Fig. S3, Table S4). This pathway is involved in stress response in plants (Chen et al., 2021). How this expansion contributes to coenocytic life cycle remains to be determined.

Overall, the global survey suggests that *Bryopsis* in essence possesses a similar set of genetic pathways to other green algal species.

### Massive duplication of genes encoding Bryohealin, a lectin required for cytoplasmic aggregation, specifically in *Bryopsis*

Next, we aimed to identify the specific genes (or gene families) that might characterise *Bryopsis*.

The best-known feature of *Bryopsis* is its amazing regeneration ability, which appears to be specific to this genus. We therefore focused on lectin, which facilitates cytoplasmic aggregation during regeneration (Kim et al., 2006; Niu et al., 2009). We searched for *BPL* lectin genes in the *Bryopsis* sp. genome and identified 12 genes highly homologous to *BPL-1* (named *BPL-1A* – *BPL-1L*) (Fig. 3). BPL-1 is characterised by a conserved ‘F-type domain’, which is widely observed in the genome of animals but not of land plants. Interestingly, the F-type domain was hardly found in other green algae genome we surveyed, and could not be identified also in *C. lentillifera* or *O. quekettii*, which belongs to Bryopsidales; we found them only in Volvocaceae among 26 green algal species surveyed in this study (Fig. 3A, Table S4). Thus, this type of lectin was lost in the majority of the green plant lineage, but dramatically expanded in *Bryopsis*.

**Figure 3.**
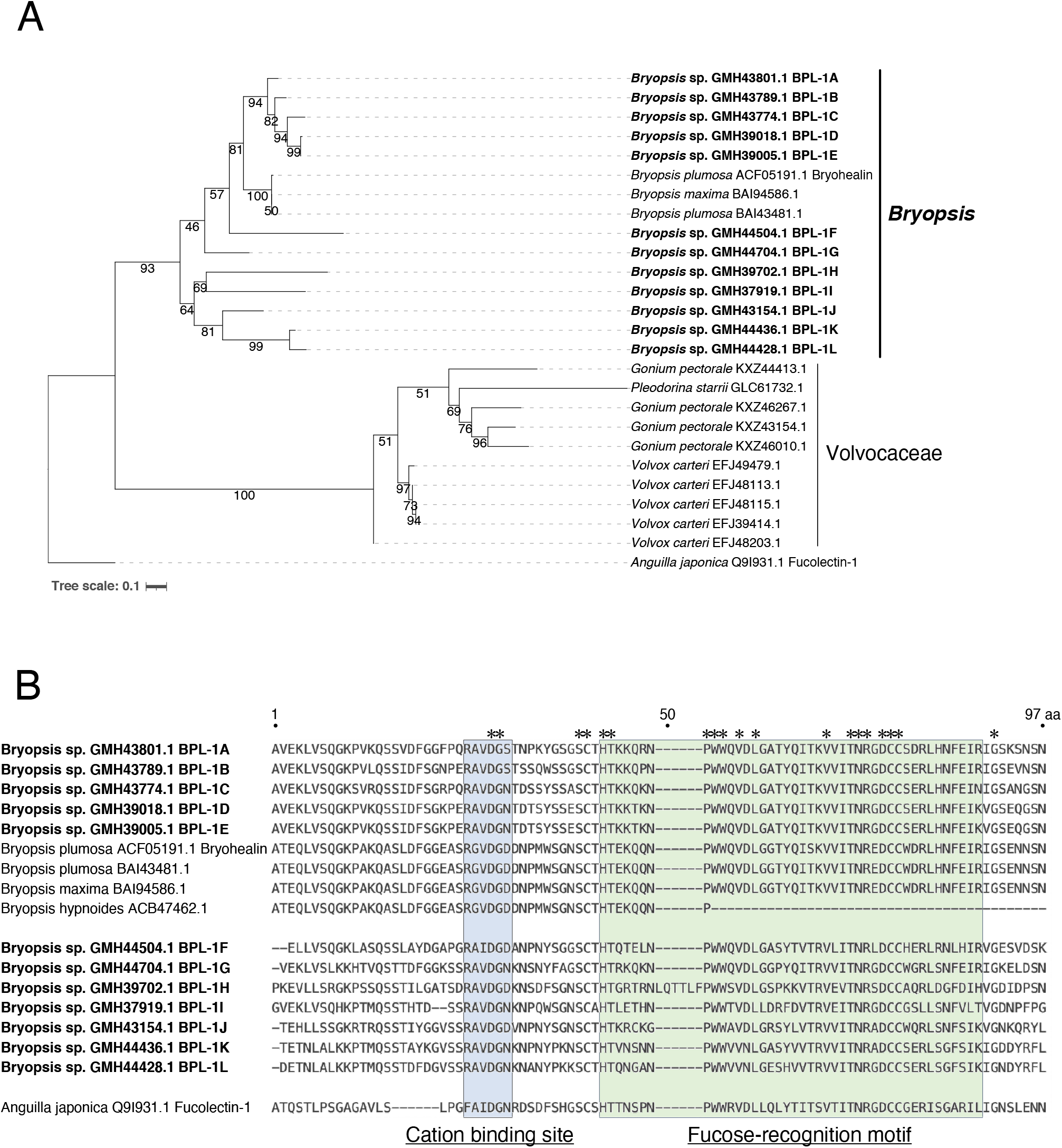
Massive duplication of BPL-1/Bryohealin in *Bryopsis* sp. (A) Phylogenetic tree of BPL-1 proteins in green algae. Only partial sequences were available for *Bryopsis hypnoides* ACB47462.1 and *Pleodorina starrii* GCL49965.1, and therefore these were not included in the tree. ML tree was constructed with WAG+G4 selected as the best-fit model and the branch support was estimated with 1,000 ultrafast bootstrap. The bar indicates 0.1 amino acids substitutions per site. (B) Alignment of amino acid sequences of BPL-1/Bryohealin of *Bryopsis* species. Asterisks indicate highly conserved residues.

BPL-2 lectin protein was also found only in *Bryopsis* (Fig. S4, Table S4). BPL-3 and BPL-4 possess the H-type domain. Our survey identified three and two homologues in the genome of *Bryopsis* sp., respectively. Unlike BPL-1 (F-type domain-containing), the H-type domain was found in the genome of *C. lentillifera* (11 genes). However, we could not identify this type of lectin in other green algae (Fig. S4, Table S4).

We searched for other lectin families, including R-type, L-type, and B-type lectins that are found in land plants, and C-type lectin and galectin that have been extensively studied in animals (Varki et al., 2022). However, we could not identify any of them. The only lectin we found was calnexin/chitinase, which is commonly present in eukaryotes.

Thus, our analysis revealed an intriguing correlation in which key lectin genes that facilitate cytoplasmic aggregation are expanded in *Bryopsis*. Lectin gene duplication might endow *Bryopsis* with its exceptional regeneration ability.

### No peculiarity in gene superfamily involved in membrane trafficking, including those essential for plant cytokinesis, in *Bryopsis*

Conceivably, the development of an extremely large cell is accompanied by a sophisticated organisation of the cytoplasm. Genes involved in membrane trafficking, which is required for cellular organisation and cytokinesis, are possibly increased or decreased in Bryopsidales.

Conserved gene families regulating membrane trafficking include the Rab GTPase, which is crucial for vesicle trafficking, and SNARE, which is required for the final step in vesicular trafficking, namely membrane fusion (Lipka et al., 2007). Previous study suggested that the increase in the number of SNARE genes parallels the rise of multicellularity among the green plants (Viridiplantae) and also Opisthokonta, based on the genome-wide survey of model species, such as *A. thaliana*, *P. patens*, *C. reinhardtii*, *Ostreococcus tauri*, *Saccharomyces cerevisiae*, and *Homo sapiens* (Sanderfoot, 2007). Similarly, the number of Rab GTPase is dramatically increased in land plants and animals compared to unicellular yeast, leading to the notion that multicellular organisms have more complex systems of internal membranous organelles than unicellular organisms (Saito and Ueda, 2009). Notably, land plants harbour a large number of Rabs and SNAREs that diverge in a manner unique to plant lineage (Saito and Ueda, 2009).

We searched for genes encoding Rab GTPase and SNARE based on BLAST and confirmed their massive increases in land plants compared to *Chlamydomonas* (Table S4). However, further survey in coenocytic Bryopsidales (*Bryopsis* sp., *C. lentillifera*, *O. quekettii*) and multicellular *Ulva*, and *Chara* (closest relative of land plants) indicated that the gene number was comparable to *Chlamydomonas*, regardless of the body form.

Among SNARE genes, *KNOLLE* is specifically required for the final step of cytokinesis, namely vesicle fusion to the cell plate; the loss of KNOLLE proteins produces multinucleated cells in land plant cells (Lauber et al., 1997; Saito and Ueda, 2009). However, this type of SNARE was present in Bryopsidales (Table S4). These results suggest that the lack of cytokinesis in *Bryopsis*’s main axis cannot be attributed to the lack of vesicle trafficking machinery.

### Cytoskeletal motor toolbox

Cytoskeleton and the associated motor proteins, which are categorised into ‘cell motility’ in KEGG database, are also key elements to cellular organisation. Microtubules and actin filaments serve as tracks for motor proteins (kinesin/dynein and myosin, respectively) to carry various cargo such as organelles. Although α/μ-tubulin and G-actin, the building blocks of microtubules and actin filaments, respectively, are highly conserved molecules, different organisms have remarkably different motor repertoires (Reddy and Day, 2001; Vale, 2003). The motor repertoire reflects the cellular dynamics and lifecycle of a species. For example, the development and function of sperm flagella requires the dynein motor as the force generator and driver of intraflagellar transport, and the loss of flagellated sperm during plant evolution coincides with the loss of dynein genes (Lucas and Geisler, 2022). Long-range transport in filamentous fungi is driven by fast and processive motor Kinesin-3, which is lost in short budding yeast (Siddiqui and Straube, 2017). Spatial distribution of mRNA encoding motor proteins may also be indicative of spatially regulated cellular activity (Andresen et al., 2021b).

We analysed cytoskeletal motor proteins based on the conserved motor domains of myosin, dynein heavy chain (DHC), and kinesin. The targeted genome sequences were of two land plant and nine green algal species (Table S2). In addition, we obtained the raw data on RNA-seq from the database for three species from Dasycladales (*Acetabularia acetabulum*, *Polyphysa clavata*, *Chlorocladus australasicus*), and two from Cladophorales (*Chlorocladiella pisiformis* and *Chlorocladiella medogensis*) (Andresen et al., 2021b; Hou et al., 2022). We assembled those sequences and annotated the genes (BUSCO values in Table S5). Dasycladales has a unique life cycle, in which a giant cytoplasm develops without nuclear division. Cladophorales is multicellular but each cell has multiple nuclei; cytokinesis is not coupled with nuclear division (Del Cortona et al., 2020; Shirae-Kurabayashi et al., 2022). For some motors, BLAST search was conducted for those species.

### Myosin

Three classes of myosin have been identified in green plants. Myosin-XI drives cytoplasmic streaming and organelle/vesicle transport in *Arabidopsis* and moss (Tamura et al., 2013; Vidali et al., 2010). Closely-related Myosin-XIII is also likely involved in intracellular transport as well as cell growth in green algae, based on localisation study in *Acetabularia* (Andresen et al., 2021b; Vugrek et al., 2003). Cytoplasmic streaming is dependent on actin filaments in the extremely large cytoplasm of *Acetabularia* (Nagai and Fukui, 1981). Myosin-VIII regulates microtubule-actin crosslinking and is required for cell tip growth, branching, and cytokinesis in moss (Wu and Bezanilla, 2014; Wu and Bezanilla, 2018; Wu et al., 2011). We anticipated that myosin genes would be conserved and the numbers possibly increased in organisms with giant cytoplasm.

This was indeed the case for Dasycladales: we identified at least five Myosin-XI/XIII in all three species examined. In surprising contrast, we identified only one myosin gene (Myosin-XI) in *Bryopsis* sp. (Fig. 4A, S5, Table S4). Other Bryopsidales species had two Myosin-XI genes, but no Myosin-VIII or –XIII. This contrasted with Cladophorales, which had multiple Myosin-XI and Myosin-XIII genes, or *U. mutabilis* and *C. reinhardtii*, where Myosin-VIII was present (Fig. 4A, B, S5, Table S4).

**Figure 4.**
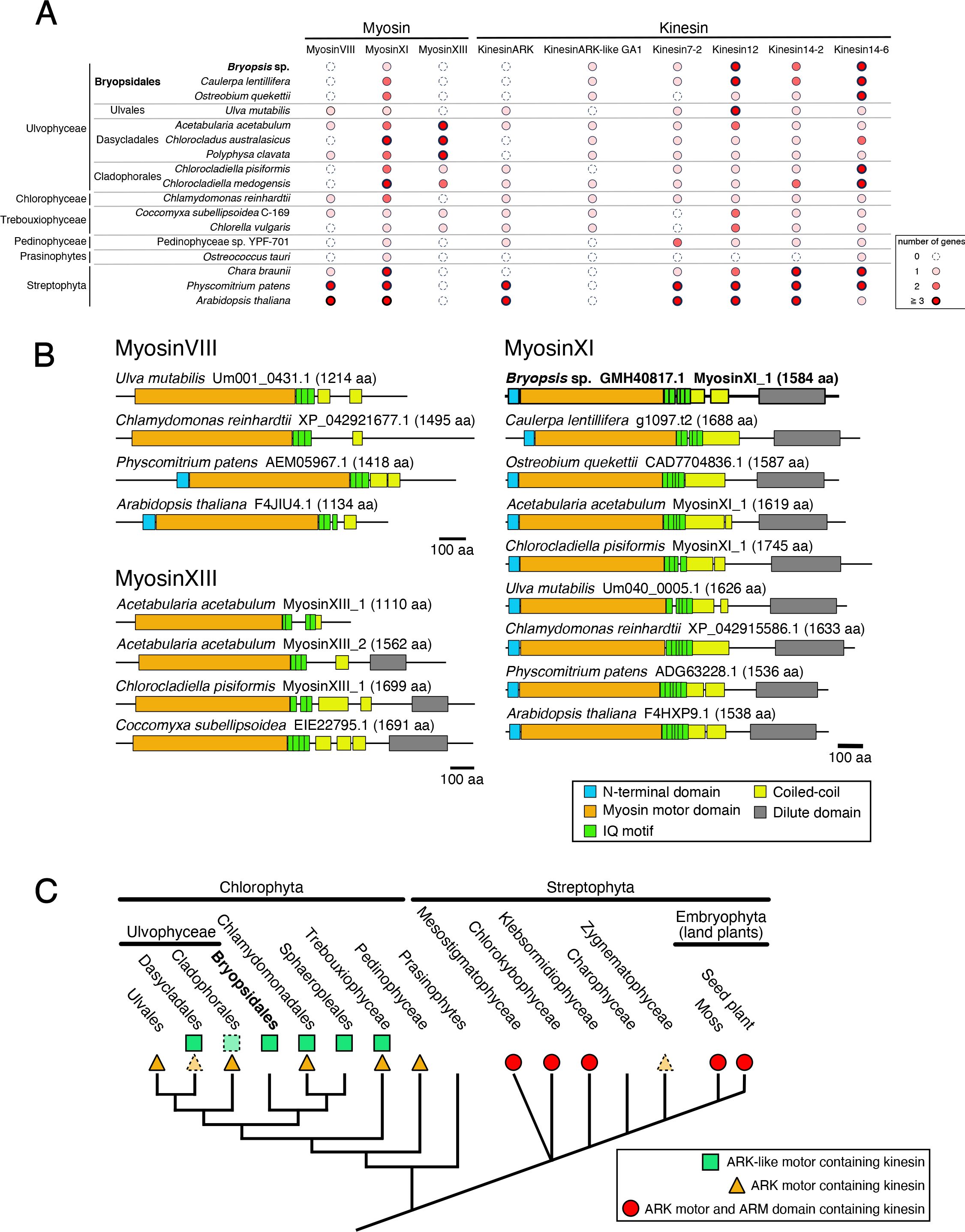
Myosin and kinesin motors in *Bryopsis* sp. (A) Repertoire of motors potentially involved in cargo transport and cytokinesis. Note that the number might be underestimated in some species, as the genome (RNA) coverage is not complete. (B) Schematic presentation of myosin motors. (C) Divergence of ARK-type motors in green plants. In case some species possess the motor but others in the same family do not, dotted lines were used.

**Figure 5.**
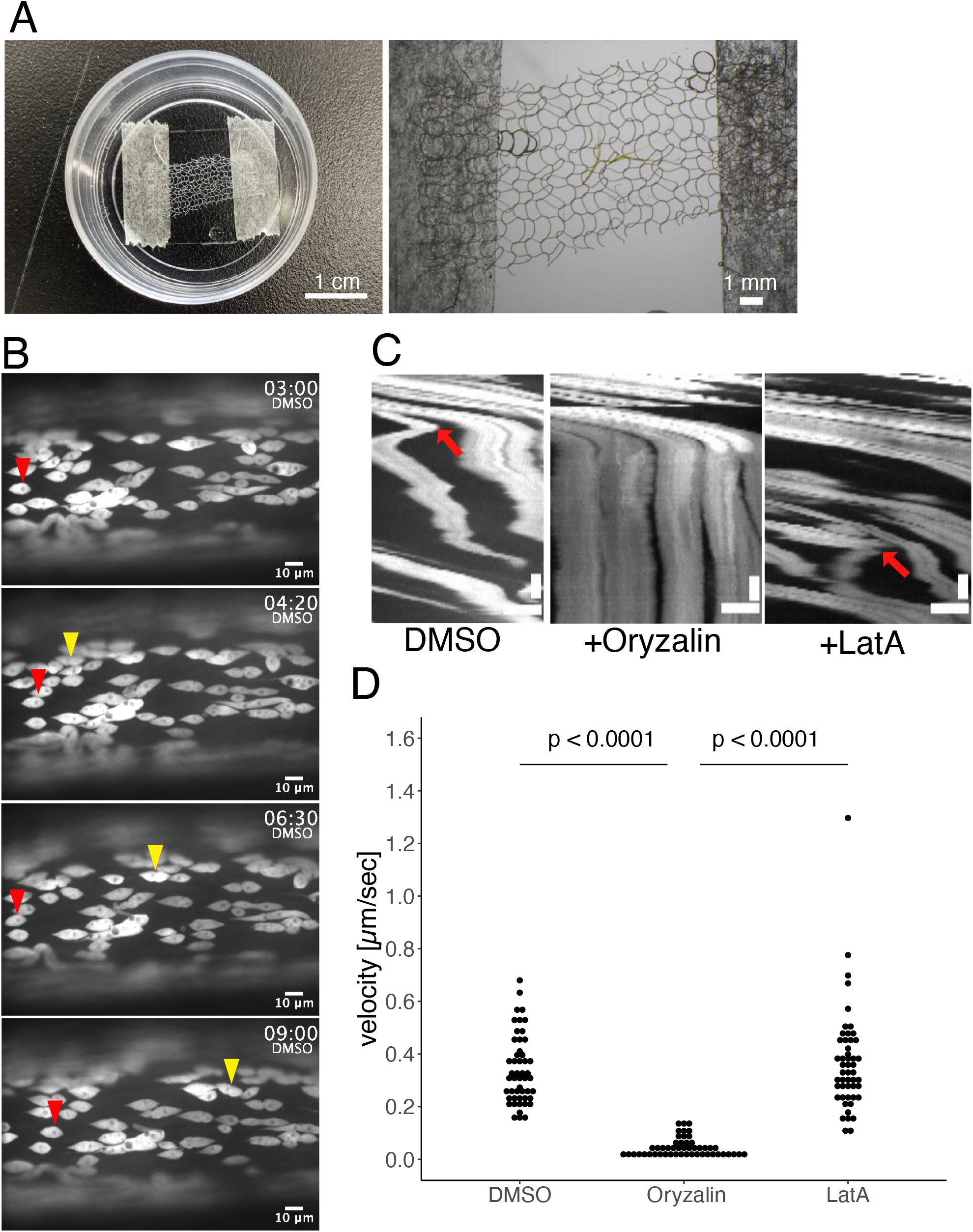
Microtubule-dependent, but actin-independent, bidirectional motility of chloroplasts in *Bryopsis* sp. (A) (Left) Device used for time-lapse imaging. (Right) Magnified view of the specimen (thalli) and a piece of net on the glass. (B) Time-lapse imaging of autofluorescent chloroplasts in the control DMSO-treated cell. Yellow and red arrowheads indicate unidirectional and bidirectional movement, respectively. Time is shown as min:sec. (C) Kymograph images of chloroplast motility in the presence or absence of microtubules or actin. Arrow indicates a point of directional switch. Horizontal bar, 10 µm; vertical bar, 120 s. (D) Rate of chloroplast motility. The mean rate was 339 ± 18 nm/s (control DMSO, ± SEM, *n* = 50), 45 ± 5 nm/s (+ oryzalin, ± SEM, *n* = 50), 369 ± 28 nm/s (+ latrunculin A [LatA], ± SEM, *n* = 50). P-values were calculated using a two-sided ART ANOVA; P < 0.0001 (control [DMSO] – oryzalin), P < 0.0001 (oryzalin – latrunculin A), P = 0.7790 (control [DMSO] – latrunculin A).

The lack of Myosin VIII in Bryopsidales and Cladophorales might be consistent with the lack of nuclear division-coupled cytokinesis in these organisms. In contrast, the underdevelopment of Myosin-XI/XIII suggests that actomyosin system is unexpectedly not prevalent in the intracellular transport of Bryopsidales.

### Dynein

Dynein is the major minus-end-directed (or ‘retrograde’) transporter in many species, except for seed plants, which lack dynein genes. Our analysis identified 13 dynein heavy chain (*DHC*) genes in *Bryopsis* sp. (Table S4). Each belongs to one of the 16 subfamilies of *C. reinhardtii* DHC (Hom et al., 2011), which consists of either the inner arm, outer arm, or intraflagellar transport (IFT) dynein complex. This was an expected finding, as flagella were present in the gametes and zoospores of *Bryopsis* sp. (Fig. 1A). We analysed the expression level of *DHC* genes based on RNA-seq. We observed that the expression of each *DHC* gene was extremely low in the main axis or rhizoid and elevated in the side branch where flagella were later developed (Table S6, p < 0.05 for 8 out of 13 genes, Likelihood ratio test). Similar *DHC* repertoire was identified in other green algal species (some genes were not identifiable either because they are absent or genome assembly is incomplete).

In the Opisthokonta lineage, ‘cytoplasmic dynein’ was evolved and acts as the major retrograde transporter in the cytoplasm of animal and fungal cells. However, we could not see the development of new types of dynein (i.e. non-flagellar dynein) in any green algal species, including *Bryopsis* sp.

### Kinesin

The kinesin superfamily has been further classified into 14 subfamilies (Lawrence et al., 2004; Shen et al., 2012). We identified a total of 34 kinesin genes in *Bryopsis* (Fig. S6.1–6.3, Table S4). Several notable features are as follows:

#### Kinesin-GA

The phylogenetic tree indicated that 20 genes belong to the canonical kinesin subfamily. Their functions can be deduced from the rich research history on kinesins in animal and plant models. However, 14 kinesins form clades that are apparently <u>g</u>reen <u>a</u>lgae-specific and do not contain plant kinesins (termed GA1–10 clades). GAs represent 40% of the total kinesins of *Bryopsis* sp.; the function of each kinesin-GA is unknown. We suggest the addition of these new subfamilies to the kinesin superfamily.

#### Kinesin-14

Land plants duplicated Kinesin-14 genes and utilise them as retrograde transporters. In *P. patens*, Kinesin-14II (KCH) is responsible for nuclear migration, whereas Kinesin-14VI (KCBP) transports the chloroplasts and others (Yamada and Goshima, 2018; Yamada et al., 2017; Yoshida et al., 2019). We identified in *Bryopsis* sp. two Kinesin-14II and three Kinesin-14VI genes, which may act as transporters (Fig. 4A, Table S4); the expression level of Kinesin-14VI is high (Table S6). Three or more Kinesin-14VI genes were found in Bryopsidales and Cladophorales, whereas Dasycladales and *Ulva* have one or two. The increase in kinesin-14VI genes and their high expression are consistent with the notion that Bryopsidales heavily utilises a microtubule-based system for cargo transport.

#### Kinesin-ARK

Animals use Kinesin-1 (also called ‘conventional kinesin’) as the versatile plus-end-directed (or ‘anterograde’) transporter, whereas ARK kinesin has recently been identified as the plant counterpart (Kanda et al., 2023; Yoshida et al., 2023). Some algal species possess a kinesin whose motor domain is similar to ARK but lacks their characteristic tail (here termed Kinesin-ARK). These are candidate anterograde transporters. However, the orthologous genes are missing in Bryopsisdales. Instead, they encode an algae-specific kinesin (kinesin-GA1) that is phylogenetically close to Kinesin-ARK (Fig. 4A, C). This kinesin subfamily possibly participates in anterograde transport; however, our RNA-seq analysis suggested that the expression level of GA1 was extremely low throughout the haploid thallus (Table S6). Therefore, it remains unclear which genes drive anterograde transport in *Bryopsis*. Intriguingly, an algae-specific Kinesin-GA9 gene (GMH32198.1) showed the highest expression level among cytoskeletal motors throughout the thallus, comparable to a sum of three Kinesin-14VIs (Table S6: total reads of this GA9 and 14VI were 535 and 498 [Deseq2]). We speculate that this novel kinesin subfamily plays an important role in *Bryopsis*, possibly as anterograde transporters.

#### Kinesin-12

Kinesin-12 genes are expanded in plants; six and 18 genes have been identified in the genomes of *A. thaliana* and *P. patens*, respectively (Shen et al., 2012). The majority of plant Kinesin-12 genes studied thus far are involved in cytokinesis. For example, plant Kinesin-12II (PAKRP) is localised in the midzone of phragmoplasts (a microtubule-based apparatus assembled in late mitosis) and is required for cytokinesis (Lee et al., 2007). Kinesin-12I (POK) is essential for the directed expansion of phragmoplasts and for division plane orientation (Livanos and Muller, 2019). In our survey, Kinesin-12II was found only in *Chara braunii* and land plants. This coincides with the development of phragmoplasts in plant evolution (Buschmann and Zachgo, 2016). However, multiple other Kinesin-12 genes, including POK-like kinesin and unclassified ones, were present in coenocytic *Bryopsis* sp. or *C. lentillifera* (Fig. 4A). They were highly expressed throughout the haploid thallus (Table S6). The result suggests that Kinesin-12I has a hitherto unknown, non-cytokinetic function in cells.

#### Kinesin-7

Mutants of Kinesin-7II (also known as NACK) fail to form the cell plate, resulting in multinucleate cells in tobacco and *Arabidopsis* (Nishihama et al., 2002; Tanaka et al., 2004). Upon sister chromatid separation in mitotic anaphase, Kinesin-7II recruits MAP kinase to the phragmoplast, by which conserved microtubule-binding protein MAP65 is phosphorylated (Sasabe and Machida, 2012). MAP65 then recruits proteins involved in vesicle trafficking for cell plate formation (Steiner et al., 2016). Thus, this kinesin acts at cytokinesis initiation. In this context, the presence of kinesin-7II in *Bryopsis* and *C. lentillifera* was unexpected (Fig. 4A). However, gamete formation in the side branch involves cellularisation in *Bryopsis*. RNA-seq analysis indicated that kinesin-7II is hardly expressed in the main axis (1.5 ± 0.49 reads, ±SD, n = 4, normalised by DESeq2) or rhizoid (0.0 ± 0.0) but is expressed at higher levels in the side branch (7.7 ± 2.6). Thus, it is tempting to speculate that the lack of cell separation in the cytoplasm in the *Bryopsis* is partly attributed to the reduced presence of this kinesin protein.

### Chloroplast motility depends on cytoplasmic microtubules, but not actin filaments

Cytoplasmic streaming in the giant cytoplasm of *Acetabularia* or in the internodal cell of *Chara* is inhibited by actin filament disassembly (Nagai and Fukui, 1981; Nagai and Kamiya, 1977). Consistent with this, multiple myosin-XIs, one of which is the fastest cytoskeletal motor (Haraguchi et al., 2022), are encoded by *C. braunii* (Fig. 4A, Table S4). Similarly, the addition of an actin polymerisation inhibitor suppressed chloroplast motility in the *Bryopsis* thallus (Menzel and Schliwa, 1986b). However, this observation was hard to reconcile with the genomics data where only one myosin gene was identified in *Bryopsis*. Therefore, we empirically revisited the contribution of microtubules and actin in intracellular transport (Fig. 5).

We focused on chloroplasts because they are autofluorescent and can be traced unambiguously using live confocal imaging. A previous study indicated that motility is dependent on both actin filaments and microtubules (Menzel and Schliwa, 1986b). We observed that chloroplasts moved along the long axis at 339 ± 131 nm/s (± SD, n =50). The movement was bidirectional and a directional switch was occasionally observed (Fig. 5B, red arrowhead; 5C, arrow; Movie 2). Motility was dependent on microtubules; oryzalin treatment almost completely abolished motility (Fig. 5C, D). Surprisingly, motility was not affected by latrunculin A treatment, although the concentration and incubation time were identical to those used when actin disappearance was confirmed by immunofluorescence microscopy (Fig. 5C, D, S1E). We presumed that cytochalasin D, which was used in a previous study to disrupt the actin cytoskeleton, has an off-target effect in *Bryopsis*. The presence of only one myosin in *Bryopsis* sp. is consistent with the notion that bidirectional transport is not driven by actomyosin. We conclude that chloroplast motility is dependent on microtubules, but not on actin filaments. The bidirectional nature of motility suggests the involvement of both retrograde and anterograde transporters. Multiplicated kinesin-14VI genes are strong candidates responsible for retrograde motility.

## Conclusions

This study provides the first information on the nuclear genome of the family Bryopsidaceae. Small contig numbers (27) and the detection of probable telomere sequences at both ends of the five contigs suggested a high-level assembly. These sequences allowed comparative genomic analyses, as illustrated here for several gene families. In addition, specialised chromosomal DNA sequences such as centromeres may be analysable. Male and female lines have been cultured in the laboratory for a few years and could, therefore, be excellent targets for developing tools for genetics and the cell and developmental biology of *Bryopsis*.

## Materials and methods

### *Bryopsis* isolation and culture

Two *Bryopsis*-like macroalgal thalli were collected on 7^th^ November 2019 from an outside tank at the Sugashima Marine Biological Laboratory. In addition to having relevant morphology and life cycle, they were confirmed to be *Bryopsis* by PCR, using primers designed for the rDNA ITS region (Shirae-Kurabayashi et al., 2022). Daily cultivation of haploid thalli was conducted at 15 °C (90 µmol m^-2^s^-1^, light: 16 h, dark: 8 h) in ocean surface water (salt concentration 2.8–3.4%), which was filtered using a 0.22-μm Millipore Stericup, autoclaved, and supplied with Daigo’s IMK medium (252 mg/L, Shiotani M.S.). Male and female gametes were obtained by culturing severed haploid thalli for 1–2 weeks at 15 °C (90 µmol m^-2^s^-1^, light: 16 h, dark: 8 h). They were mixed and cultured under the same conditions for ∼1 week. Once sporophyte (diploid) germination was detected, the culture condition was changed (25 °C, 20 µmol m^-2^s^-1^, light: 10 h, dark: 14 h). After six months, the cells darkened. Culture conditions were changed (15 °C, 90 µmol m^-2^s^-1^, light: 16 h, dark: 8 h). Zoospores (haploids) were released under these conditions, followed by germination in ∼1 week.

### Protoplast formation from extruded cytoplasm

The thallus was cut with a scalpel, and sandwiched and crushed with two slide glasses. The extruded cytoplasm was slowly dripped into autoclaved seawater in the presence or absence of N-acetyl-D-glucosamine (40 mM) or the control glucose (40 mM).

### RNA sequencing (RNA-seq)

#### For genome assembly and gene annotation

*The Bryopsis* sample (female line, *Bryopsis* sp. KO-2023) was crushed in liquid nitrogen, and the total RNA was purified using the RNeasy Plant Mini Kit (#74904; Qiagen, Hilden, Germany) with DNase treatment, according to the manufacturer’s instructions. The RNA yield was quantified using a NanoVue microplate reader (GE Healthcare, Chicago, IL, USA). The sample volume was adjusted to 2 µg/100 µL for subsequent RNA-seq analysis. RNA-seq analysis was performed at the core facility of Nagoya University following the protocol described by (Matsumura et al., 2022). Briefly, 1 µg total RNA was used for mRNA purification with NEBNext Oligo d(T)_25_ (NEBNext poly(A) mRNA Magnetic Isolation Module; New England Biolabs, Ipswich, MA, USA), followed by first-strand cDNA synthesis with the NEBNext Ultra II RNA Library Prep Kit for Illumina (New England Biolabs) and NEBNext Multiplex Oligo for Illumina (New England Biolabs) according to the manufacturer’s protocols. The amount of cDNA was determined using an Agilent 4150 TapeStation System (Agilent, Santa Clara, CA, USA). The cDNA libraries were sequenced as paired-end reads of 81 nucleotides using an Illumina NextSeq 550 (Illumina, San Diego, CA, USA).

#### Spatial dissection

Fragments of < 1 mm from the tip of the main axis of *Bryopsis* sp. were cut and cultured in autoclaved seawater supplemented with Daigo’s IMK medium for 10 –14 days at 15 °C (90 µmol m^-2^s^-1^, light: 16 h, dark: 8 h). The thalli that developed side branches were cut into three parts; ‘side branch’, ‘main axis’ (central stalk), and ‘rhizoid’. After removing water, each sample was separately crushed with mortar and pestle that had been prechilled at –80 °C, and the total RNA was purified using the RNeasy Plant Mini Kit. This manipulation was independently performed four times on different days. RIN values for all samples were greater than 8.0. The samples were sequenced with Illumina NovaSeq6000 platform, which produced 150 bp paired-end reads. The amount of reads for each gene was calculated using RSEM v1.2.28 (Li and Dewey, 2011) with STAR v2.7.10b (Dobin et al., 2012) for mapping. Normalisation was performed using TPM and DESeq2 (Love et al., 2014).

### Genome sequencing

Whole-genome shotgun sequencing was performed using the PacBio and Illumina sequencing platforms. Genomic DNA from *Bryopsis* sp. KO-2023 (female) was isolated using a CTAB/Genomic-tip Kit (QIAGEN). A SMRTbell library for continuous long-read (CLR) sequencing was prepared using a SMRTbell Express Template Prep Kit 2.0 (Pacific Bioscience, CA, USA) according to the manufacturer’s instructions. The CLR library was size-selected using the BluePippin system (Sage Science, Beverly, MA, USA) with a lower cutoff of 30 kb. One SMRT Cell 8M was sequenced on the PacBio Sequel II system with Binding Kit 2.0 and Sequencing Kit 2.0 (20 h collection times). In addition, genomic DNA was fragmented to an average size of 500 bp using an M220 Focused-ultrasonicator M220 (Covaris Inc., Woburn, MA. USA). A paired-end library with insert sizes ranging from 450 to 550 bp was constructed using the TruSeq DNA PCR-Free Library Prep kit (Illumina) and was size-selected on an agarose gel using a Zymoclean Large Fragment DNA Recovery Kit (Zymo Research, Irvine, CA. USA). The final library was sequenced using a 2 × 150 bp paired-end protocol on the NovaSeq 6000 system (Illumina).

### Genome assembly

#### Chloroplast

*De novo* assembly of the chloroplast genome was performed using a combination of 150 bp × 2 short reads and Get-organelle v 1.7.6.1 (Jin et al., 2020) with the options –k 21, 45, 65, 85, 105, –P 1000000, and –R 50. Two complete *Bryopsis* chloroplast sequences (NC_026795.1 and NC_013359.1) were used as seeds. This provided two closed circular sequences of identical length (91,672 nt). The two sequences were nearly identical except for the central region (∼11 kb). One sequence was discarded because structural errors were found near the central region when it was aligned with long reads. The other sequences showed no structural errors across the entire sequence length. The error check was repeated at different starting positions. Finally, the downstream of *psbA* was set at +1 position.

#### Mitochondrion

Highly fragmented contigs with a total length of ∼150 kb were obtained using Get-organelle v 1.7.6.1 assembly (Jin et al., 2020) with the seed references of green algal species (NC_045361.1, KU161104.1, and NC_001638.1) (Repetti et al., 2020; Vahrenholz et al., 1993; Zhou et al., 2016). These putative mitochondrial sequences had a sequencing depth ∼200 times higher than that of the nuclear genome. The high copy number of the mitochondrial genome enabled assembly based on random selection of a small portion of PacBio long reads (≥20 kb). One percent of the long reads was sufficient for the assembly of the mitochondrial genome. Flye (Kolmogorov et al., 2019; Lin et al., 2016) with ‘--pacbio-raw’ option produced one circular sequence (356,161 bp) that had global synteny with other algal mitochondrial sequences. To check if there was mis-assembly in this sequence, full long and short reads were aligned using minimap2 (Li, 2021) with the ‘map-pb’ and ‘sr’ presets, respectively. This revealed six indel errors at the homopolymer sites but did not identify any large sequence gaps or structural errors. Small indels were corrected using bwa (mapping) and Pilon (Walker et al., 2014). To confirm the completeness of the mitochondrial genome assembly, the +1 position was changed by 20,000 bp and the long reads were aligned using minimap2. No sequence gaps were found during this operation, indicating that no structural errors existed in the mitochondrial assembly. Finally, the +1 position was reset downstream of *rrnL3b*.

#### Nuclear genome

The assembly of long-read data was used to determine the nuclear genome. However, the genome sequences of symbiotic bacteria, commonly detected in marine macroalgae, inevitably contaminate *Bryopsis* genome sequences. Therefore, a provisional genome assembly was first performed, in which the obtained genome sequences were clustered into groups which were thought to originate from the same species. Based on the sequence characteristics and mapping results of the RNA-seq data, grouped sequences considered to be derived from *Bryopsis* were identified. Sequences were extracted from clustered groups.

Illumina reads were used for K-mer analysis and genome size estimation. The 21-mer frequencies were calculated using Jellyfish v2.3.0 (Marcais and Kingsford, 2011), and the genome size was estimated using GenomeScope 2.0 (Ranallo-Benavidez et al., 2020). The estimated genome size was used as the input parameter for *de novo* pre-assembly. Pre-*de novo* assembly of the nuclear genome was performed based on the PacBio reads using Canu v2.1.1 (Koren et al., 2017) with the following options: genomeSize = 500M, corOutCoverage = 200, and ‘batOptions = –dg 3 –db 3 –dr 1 –ca 500 –cp 50’. Pre-assembled contigs were polished using long and short reads. They were polished through three rounds of Arrow v2.3.3, and three rounds of Pilon v1.23 (Walker et al., 2014). In these steps, PacBio reads were mapped using pbmm2 v1.3.0 (https://github.com/PacificBiosciences/pbmm2), and trimmed Illumina reads were mapped using BWA v0.7.17 (Li, 2013). Then, binning was performed using MetaBAT2 v2.15 (Kang et al., 2019) to group contigs derived from the same species, and each cluster was named ‘bin’. As input for MetaBat2, read coverage information was calculated from the Illumina read mapping results against polished pre-assembled contigs using BWA v0.7.17.

Raw RNA-seq data were trimmed and filtered using Platanus_trim v1.0.7. *De novo* transcriptome assembly was performed based on the trimmed RNA-seq reads using Trinity v2.8.5 (Grabherr et al., 2011). Transcriptome assembly contigs were splice-mapped to polished, pre-assembled genomic contigs using GMAP v.2018-08-25 (Wu and Watanabe, 2005). The bin containing the most-mapped transcriptome assembly contigs was designated as the main nuclear bin. In addition, other bins and contigs derived from *Bryopsis* were manually selected based on the overall information, such as the transcriptome assembly contig mapping rate, GC rate, and Illumina read coverage.

PacBio and Illumina reads derived from *Bryopsis* were extracted for the final *de novo* assembly. PacBio reads were extracted from Canu intermediate files used in the pre-*de novo* assembly. Illumina reads were extracted by mapping the trimmed Illumina reads to contigs derived from *Bryopsis* using BWA v0.7.17. The extracted trimmed Illumina reads were used for K-mer analysis and genome size estimation, as described above. The estimated genome size was used as an input parameter for the final *de novo* assembly. Final *de novo* assembly of the nuclear genome was performed based on the PacBio reads derived from *Bryopsis* using Canu v2.2 with the following options: genomeSize = 100M, corOutCoverage = 200, and ‘batOptions= –dg 3 –db 3 –dr 1 –ca 500 –cp 50’. The final assembled contigs were polished using long and short reads. The final assembly contigs were polished through three rounds of Arrow v2.3.3 and three rounds of NextPolish v1.4.0 (Hu et al., 2019). Next, the arrow-identified variants were filtered via Merfin v1.0 (Formenti et al., 2022) using the trimmed Illumina reads derived from *Bryopsis*. In the long-read-based polish, PacBio reads derived from *Bryopsis* were mapped using pbmm2 v1.3.0. Haplotigs were then removed using Purge_dups v1.2.3 (Guan et al., 2020) to reduce sequence redundancy and increase assembly continuity.

These analyses yielded the assembly and selection of 49 contigs. Finally, to verify the origin of each contig, BLASTx searches were conducted for a portion of the sequence of each contig. The sequences derived from 22 contigs were highly homologous to bacterial and fungal sequences, whereas those of the other 27 contigs were not. Thus, 27 contigs were considered derived from *Bryopsis*.

### Gene annotation

#### Chloroplast

ncRNAs were annotated using the GeSeq web server. ‘DNA search identity’ was set at 85. Four reference sequences (NC_013359.1, NC_026795.1, NC_037363.1, and NC_ 030629.1) were used as ‘3rd Party References.’ The CDS was manually annotated using a combination of GeSeq annotation, protein alignment with *B. plumosa* (NC_026795.1), and RNA-seq alignment. This collaborative annotation was further curated using a homology-based approach against the proteomes of closely related species to verify the completeness of each CDS. In total, 83 predicted protein-coding genes, three rRNAs, and 26 tRNAs were identified.

#### Mitochondrion

Annotation of the mitochondrial genome using GeSeq predicted virtually no protein-coding genes. This suggests that no closely related protein-coding genes were annotated. To overcome this limitation, open reading frames (ORFs) were searched using the NCBI ORF finder (https://www. ncbi.nlm.nih.gov/orffinder/). The predicted ORFs of all six frames were manually aligned with the mitochondrial proteins of *Ostreobium quekettii* (Repetti et al., 2020) and the putative CDS coding frame of the RNA-seq was constructed with TransDecorder. To verify the obtained CDS, promising coding frames were manually searched for homology to proteins of closely related species using BLASTx. This procedure identified 40 protein-coding genes with complete CDS sequences. In addition, a tBLASTn search using publicly available green algal mitochondrial protein sequences identified 14 small genes encoded in the introns of already annotated genes. ncRNAs were annotated using the Geseq web server. tRNAs were identified with Geseq, where the following ‘3rd Party References’ were used: NC_045361.1, NC_001638.1, NC_028538.1, NC_035722.1, NC_029701.1, NC_035809.1, NC_28081.1, NC_040163.1, and NC_041082.1. This resulted in 17 annotated tRNAs. rRNAs were searched against the mitochondrial genome using BLASTn with the following queries: NC_045361.1, NC_001638.1, NC_028538.1, NC_035722.1, NC_029701.1, NC_035809.1, NC_28081.1, NC_040163.1, and NC_041082.1. Candidate genes were manually compared with the RNA-seq alignment data. This procedure identified three rRNA genes in the mitochondrial genome. Intron length is defined as the length of the region between exons within a gene (protein-coding or non-coding). When other genes were present within the introns of a host gene, the length of the internal gene was not excluded from the intron length of the host gene. Domains of genes present in introns were searched using NCBI’s Conserved Domains database (https://www.ncbi.nlm.nih.gov/Structure/cdd/wrpsb.cgi) with default settings.

#### Nucleus

Protein-coding genes were predicted by combining the results of RNA-seq-, homology-, and *ab initio*-based prediction methods. RNA-seq-based prediction utilises both assembly-first and mapping-first methods. For the assembly-first method, RNA-seq data were assembled using Trinity v2.12.0 (Grabherr et al., 2011) and Oases v2.0.9 (Schulz et al., 2012). The redundant assembled RNA contigs were removed using CD-HIT v4.8.1 (Fu et al., 2012), and then splice-mapped to the genome sequences using GMAP v2018-07-04 (Wu and Watanabe, 2005). For the mapping-first method, RNA-seq data were mapped to genome scaffolds using HISAT2 v2.2.1 (Kim et al., 2019), and gene sets were predicted with StringTie v2.2.0 (Pertea et al., 2016) from mapped results. The ORF regions were estimated using TransDecorder v5.5.0 (https://github.com/TransDecoder/TransDecoder) from both the assembly-first and mapping-first method results. Regarding homology-based prediction, amino acid sequences of *O. quekettii* (NCBI accession No: GCA_905146915.1), *C. reinhardtii* (NCBI accession No: GCF_000002595.2), *Volvox carteri* (NCBI accession No: GCF_000143455.1), and *Monoraphidium neglectum* (NCBI accession No: GCF_000611645.1), were splice-mapped to genome scaffolds using Spaln v2.3.3f (Gotoh, 2008), and gene sets were predicted. For *ab initio* prediction, training sets were first selected from the RNA-seq-based prediction results. Then, AUGUSTUS v3.3.3 (Stanke and Waack, 2003) was trained using this set. The SNAP v2006-07-28 (Korf, 2004) was used in this study. All predicted genes were combined using an in-house merging tool. However, the ORF of some genes did not start with ATG (methionine), which was manually fixed. In some cases, the start codon was manually identified, and the amino acid sequences were corrected. In other cases (∼700), the ORF assignment was rejected as the start codon and transcript could not be identified. Finally, 14,034 genes encoding proteins were identified.

### *De novo* transcriptome assembly and annotation

*De novo* transcriptome assembly and gene annotation were conducted based on the published RNA-seq raw data, following the methods described in (Andresen et al., 2021b) and (Hou et al., 2022) for the following species: [Dasycladales] *Acetabularia acetabulum*, *Chlorocladus australasicus* and *Polyphysa clavata*; [Cladophorales] *Chlorocladiella pisiformis* and *Chlorocladiella medogensis* (Supplementary Data). The raw sequence data were obtained from the European Nucleotide Archive under the accession No. PRJEB40460 and PRJNA726747.

### Genome information used in this study

The genomes primarily used in each analysis were *Bryopsis* sp. KO-2023 (this study), *C. lentillifera* (Arimoto et al., 2019), *O. quekettii* (Iha et al., 2021), *U. mutabilis* (De Clerck et al., 2018), *C. reinhardtii* (Merchant et al., 2007), *Dunaliella salina* (Polle et al., 2017), *Pleodorina starrii* (Takahashi et al., 2023), *V. carteri* (Prochnik et al., 2010), *Raphidocelis subcapitata* (Suzuki et al., 2018), *Monoraphidium neglectum* (Bogen et al., 2013), *Auxenochlorella protothecoides* (Gao et al., 2014), *Coccomyxa subellipsoidea* C-169 (Blanc et al., 2012), *Chlorella vulgaris* (Cecchin et al., 2019), Pedinophyceae sp. YPF-701 (Repetti et al., 2022), *Chloropicon primus* (GCA_023205875.1), *Micromonas pusilla* (Worden et al., 2009), *O. tauri* (Blanc-Mathieu et al., 2014), and *Bathycoccus prasinos* (Yau et al., 2020) for Chlorophyta and *Klebsormidium nitens* (Hori et al., 2014), *C. braunii* (Nishiyama et al., 2018), *P. patens* (Lang et al., 2018) and *A. thaliana* (Lin et al., 1999; Mayer et al., 1999; Salanoubat et al., 2000; Tabata et al., 2000; Theologis et al., 2000) for Streptophyta (Table S2). Note that the available *A. acetabulum* genome sequences were not amenable to comparative genomics due to low quality (BUSCO <11%) (Andresen et al., 2021a).

### Circular visualization of the genome assembly (Circos plot)

The genomic features of the 27 contigs were plotted in a circular genome plot using shinyCircus V2.0 hosted in a local server (Wang et al., 2023). GC content was calculated as the ratio of the average of AT and GC per 10,000 bp. For repetitive sequences plot, all types of repeats were used from the result of repeatmasker (see below). All information used for the circus-plot is available (https://github.com/KantaOchiai/Bryopsis_sp._KO-2023_genome_sequence_Information).

### Comparative genomics analysis

#### Repetitive sequences

Repetitive sequences were identified using a combination of *de novo* and homology-based methods. First, Repeat sequences were *de novo* searched using RepeatModeler v2.0.1 (http://www.repeatmasker.org/RepeatModeler/) with “--LTRstruct”. Then, identified repetitive sequences, including transposable elements, were counted using RepeatMasker v4.1.1 (http://www.repeatmasker.org/) based on the repeat model created by RepeatModeler (Table S1).

#### Evaluation of assembly quality

BUSCO metrics were used to assess the integrity of the genome assembly and the completeness of the gene prediction (Waterhouse et al., 2017). BUSCO v5.5.0 was run with genome or protein mode on 18 published genomes of Chlorophyta, including *Bryopsis* sp. with Chlorophyta dataset (chlorophyta_odb10), and four published genomes of Streptophyta with the Viridiplantae (viridiplantae_odb10) or Brassicales (brassicales_odb10) dataset (Table S1). The transcriptome mode was applied for transcriptomes of two Cladophorales and three Dasycladales with Chlorophyta dataset (chlorophyta_odb10) (Table S5).

#### Functional annotation with KEGG database

Functional annotation was performed based on KEGG (Kyoto Encyclopedia of Genes and Genomes) using GhostKoala (Kanehisa et al., 2016). The unigenes of each pathway in each genome were counted with KEGG mapper (https://www.genome.jp/kegg/mapper/) (Table S3). Subsequently, ‘MAPK signaling pathway-plants’ in the ‘Signal transduction’ category was analysed with BLASTp searches using the representative *A. thaliana* proteins as queries, as extremely high number of genes were identified in this category for Bryopsidales including *Bryopsis* sp. (accession No: PYR/PYL/RCARs (NP_180174.1, O49686.1, NP_563626.1), PP2C_GroupA (P49598.1), HOS15 (Q9FN19.1), RBOH (O48538.1, Q9FIJ0.1), KAT1 (Q39128.1), QUAC1 (O49696.1), SLAC1 (Q9LD83.1), ABFs/ABI (Q9M7Q3.1, Q9SJN0.1, Q9M7Q5.1), SOD (AEE74978.1, AEE85010.1), CAT1 (Q96528.3)). SnRK2 annotated with KEGG was confirmed by KEGG BLASTp web server (Fig. S3, Table S4).

### Phylogenetic inference

#### Chlorophyta species

10 highly conserved single-copy OGs were selected from 63 single copy ortholog genes (OGs) obtained using Orthofinder v2.3.14 (Emms and Kelly, 2019) in 18 published genomes of Chlorophyta including *Bryopsis* sp. and three Streptophyta (Table S2). 10 single-copy OGs list is available in Supplementary Data. Each OG sequences were aligned using MAFFT v7.505 (Katoh and Standley, 2013) with FFT-NE-2 strategy. All gaps were removed using MEGAX (Kumar et al., 2018), and the individual OGs were combined to obtain a sequence of 2,713 amino acids (Supplementary Data). Finally, ML tree was inferred using IQ-TREE v1.6.12 (Nguyen et al., 2015) with LG+F+R4 selected as the best-fit model and branch support estimated with ultrafast 1,000 bootstrap.

#### Mitochondrial genome

Seven mitochondrial housekeeping genes, including *nad1*, *nad2*, *nad4*, *nad5*, *nad6*, *cob*, *cox1*) were retrieved from 17 species, including *Bryopsis* sp. and registered *B. plumosa* (MN853874.1) (Fig. S2). The same procedure as for chloroplasts was used for the subsequent analysis.

#### Lectin

BLASTp/tBLASTn searches were conducted for published *Bryopsis* BPL-1, –2, – 3, and –4 proteins. For all possible hit sequences (Supplementary Data), the presence of characteristic domains of each BPL protein was confirmed with the NCBI conserved domain search (https://www.ncbi.nlm.nih.gov/Structure/cdd/wrpsb.cgi). BLASTp/tBLASTn searches were also conducted against *Bryopsis* sp. for R-, L-, B– and C-type lectins (accession No: P06750.1, PWZ39448.1, AAL09432.1, Q9FVA1.1, Q9FV99.1, Q9NNX6.1), malectin (accession No: AEE78805.1), calnexin (accession No: KAB1259615.1), calreticulin (accession No: CAA55890.1), chitinase (accession No: AEC10291.1), and galectin (accession No: KAJ0248405.1) as queries. Amino acid sequences of each gene were aligned by MAFFT v7.505 with FFT-NE-2 strategy. All gaps were removed using MEGAX, and sequences of 116 amino acids (BPL-1), 132 amino acids (BPL-2), and 102 amino acids (BPL-3/4) were obtained (Supplementary Data). ML tree was drawn using IQ-TREE v1.6.12 with WAG+G4 (BPL-1, –2) or LG+G4 (BPL-3/4) selected as the best-fit model and branch support was estimated with 1,000 ultrafast bootstrap.

#### Rab GTPase and SNARE

Genes were searched with BLASTp/tBLASTn using the representative *A. thaliana* proteins as queries (Rab GTPase accession No: NP_568678.1, SNARE: (Lipka et al., 2007)).

#### Myosin

Genes were searched with BLASTp/tBLASTn in nine genomes of Chlorophyta, including *Bryopsis* sp., three genomes of Streptophyta, and five transcriptomes of Cladophorales and Dasycladales (Fig. 4A, Table S2), using the following queries: Myosin-VIII (accession No: F4JIU4.1), Myosin-XI (accession No: F4HXP9.1, GMH40817.1), and Myosin-XIII (accession No: AAB53061.1, AAB53062.1). All hit sequences with the e-value ≤ e^-10^ were subjected to the NCBI conserved domain search, and the sequences in which conserved motor domains could not be identified were removed from the list (Supplementary Data). Some myosin proteins, for which long amino acid sequences could be retrieved, were shown as schematic diagrams (Fig. 4B) and/or subjected to phylogenetic tree construction (Fig. S5). For tree construction, the amino acid sequences were aligned by MAFFT v7.505 with FFT-NE-2 strategy and all gaps were removed using MEGAX, and a sequence of 184 amino acids was obtained (Supplementary Data). ML tree was drawn using IQ-TREE v1.6.12 with LG+I+G4 selected as the best-fit model and branch support was estimated with 1,000 ultrafast bootstrap (Fig. S5).

#### Dynein heavy chain (DHC)

Genes were searched with BLASTp/tBLASTn using previously reported *C. reinhardtii* DHC1–16 proteins (Hom et al., 2011) as queries (Table S4).

#### Kinesin

Genes were searched with BLASTp/tBLASTn using the amino acid sequences of 1–350 aa of the human kinesin heavy chain (KIF5B/kinesin-1: accession No: P33176.1) and *Arabidopsis thaliana* KIN4C (accession No: F4K0J3.2) as queries. Additional BLASTp/tBLASTn searches were conducted for several kinesins: Kinesin-ARK and Kinesin-GA1 (ARK-like) in 10 genomes of Chlorophyta, seven genomes of Streptophyta, and five transcriptomes of Cladophorales and Dasycladales; Kinesin-7II, Kinesin-12, Kinesin-14II, and Kinesin-14VI in five transcriptomes of Cladophorales and Dasycladales (Table S2). All hit sequences with the e-value ≤ e^-10^ were subjected to the NCBI conserved domain search, and the sequences in which conserved motor domains could not be identified were removed from the list (Supplementary Data). The kinesin amino acid sequences in nine published genomes of Chlorophyta, including *Bryopsis* sp., and three Streptophyta were aligned by MAFFT v7.505 with FFT-NE-2 strategy and all gaps were removed using MEGAX, followed by ML tree construction (IQ-TREE v1.6.12 with LG+I+G4 and branch support was estimated with 1000 ultrafast bootstrap) (Fig. S6).

### Immunostaining

A three-week-old thallus after cytoplasm extrusion was fixed with 4% paraformaldehyde in modified PHEM buffer (Sobue et al., 1988) (60 mM Pipes, 25 mM Hepes, 0.5 M NaCl, 10 mM EGTA, 2 mM MgCl_2_; pH 6.9) for 1 h at 25 °C, followed by permeabilisation with 1% Triton X-100 in PBS for 1 h at 25°C. After washing twice with PBST (0.1% Triton X-100 in PBS), the specimen was incubated with blocking solution (1% BSA in PBST) for 1 h at 25 °C, followed by addition of primary antibodies at 4 °C overnight with rotation (mouse anti-ϕ3-actin [Proteintech, 66009-1-Ig], 1:1000, and rat anti-χξ-tubulin [YOL1/34, MCA78G, Bio-Rad], 1:1000). The specimen was washed three times with PBST and incubated with secondary antibodies (anti-mouse, Jackson ImmunoResearch, 715-545-151, 1:1000, and anti-rat, Jackson ImmunoResearch, 712-165-153, 1:1000) and DAPI (final 1 µg/ml) overnight at 4 °C with rotation. After washing twice with PBST, the specimen was mounted on a glass slide with a mounting medium (Fluoromount^TM^; Diagnostic BioSystems).

### Microscopy

*Bryopsis* sp. thalli were imaged using a Nikon SMZ800N stereo microscope, Plan Apo 1×/WF lens, and NY1S-EA camera (SONY). The gametes and zoospores were imaged using an ECLIPSE E200 microscope (Nikon) and NY1-EA2. Fluorescent images of DNA (DAPI), chloroplasts, microtubules, and actin were acquired using a Nikon Ti2 inverted microscope equipped with a CSU-10 spinning-disc confocal scanner unit (Yokogawa), a Zyla CMOS camera (Andor), and four laser lines (637, 561, 488, and 405 nm). 40× 0.95 NA lens or a 100× 1.40 NA lens was used to image live or fixed cells, respectively. To obtain the chloroplast motility rate, a 35-mm glass-bottom dish was prepared, on which a piece of kitchen garbage net (∼10 × 20 mm) was attached with double-sided tape. After cytoplasmic extrusion, a 3-week-old thallus and a coverslip were laid over the net, followed by the addition of 1-mL of autoclaved seawater. This net prevented thallus movement during imaging. Autofluorescent chloroplasts were imaged every 10 s using a spinning-disc confocal microscope and a 40× 0.95 NA lens. At 2 min during imaging of untreated specimen, oryzalin (10 µM), latrunculin A (10 µM), or control DMSO was added (3 mL volume each). The unidirectional motility rate of randomly selected chloroplasts 5–6 min after drug addition was manually measured after obtaining kymograph images using Fiji.

## Data availability

The genome sequence of *Bryopsis* sp. is available at the DNA Data Bank of Japan (DDBJ) under project PRJDB15746 (https://ddbj.nig.ac.jp/resource/bioproject/PRJDB15746) and sample accession SAMD00599708 (https://ddbj.nig.ac.jp/resource/biosample/SAMD00599708) with accession numbers BSYQ01000001.1–BSYQ01000027.1 (nuclear genome), LC768901 (chloroplast), and LC768902 (mitochondria). The raw sequence data for NextSeq 550, NovaSeq 6000, and Sequel II are available under accession numbers DRA016305, DRA016314, and DRA016315, respectively. The assembled genome and annotation are also available from NCBI with GenBank accession ID: GCA_030272585.1. The IDs of the genes used for the phylogenetic tree construction are shown in the figures. Gene and protein sequences used for phylogenetic tree construction and comparative genomic analyses are summarised in Supplementary data (https://github.com/KantaOchiai/Bryopsis_sp._KO-2023_genome_sequence_Information).

## Acknowledgements

We are grateful to the staff of the Comparative Genomics Laboratory at NIG for supporting genome sequencing. This work was funded by the Japan Society for the Promotion of Science KAKENHI (16H06279 (PAGS) for whole-genome sequencing and 22K19308, 22H04717, and 22H02644 for experimental biology). The authors declare no conflict of interest.

## Supplementary document

### Overview of the chloroplast genome

In this study, the chloroplast genome was assembled into a single closed sequence of 91,672 base pairs (bp). This length was close to the size of previously reported chloroplast genomes of *Bryopsis plumosa* (106,859 bp) (Leliaert and Lopez-Bautista, 2015) and *Bryopsis hypnoides* (153,429 bp) (Lu et al., 2011). No long reverse repeat sequences were identified, consistent with other green algae of the order Bryopsidales of the family Ulvophyceae, and genus *Ulva* (Turmel and Lemieux, 2018; Turmel et al., 2017). The GC content was 30.4%, which was similar to the reported chloroplast genomes of *B. plumosa* (30.8%) (Leliaert and Lopez-Bautista, 2015) and *B. hypnoides* (33.1%) (Lu et al., 2011). The coding DNA sequences occupied 85.4% of the chloroplast genome, which was much higher than that of the mitochondria (66.1%) (Table 2, Table S7, S9). Drastic expansion of introns, which was evident in the mitochondrial genome, was not observed in either *Bryopsis* lines.

GeSeq-based annotation revealed that the chloroplast genome contained 83 protein-coding genes, 79 of which were identical to the previously annotated *bona fide* or hypothetical protein-coding genes of *B. plumosa* (NC_026795.1) and were conserved in other green algae (Table S8). The remaining four protein-coding genes included two open reading frames (ORFs) found within the introns of *psaA* and *psbB*, one previously reported ORF, and one novel ORF. The two ORFs in the introns showed high homology with the previously reported *orf1* and *orf2* of *B. plumosa* (NC_026795.1). ORF480 (i.e. 480 a.a.) in the intron of *psaA* encodes a protein that has a reverse transcriptase-like superfamily and RVT_N superfamily domains, suggesting that it functions as a reverse transcriptase. In contrast, ORF300 in the intron of *psbB* did not contain any characteristic domains, suggesting that it might not represent a protein.

One of the two isolated ORFs, termed ORF92, is a 281 bp reading frame (i.e. 92 a.a.) found in a ∼2.5 kb flanking region between *chlN* and *trnL*. RNA-seq analysis indicated that this gene was transcribed *in vivo*. However, the translated sequences showed no homology to known proteins in the database. Thus, this might be specifically encoded in the chloroplast genome of our line. The other orphan ORF, termed ORF431, showed weak sequence identity with GIIM superfamily proteins (group II intron, maturase-specific domain) according to a domain search (CD-search). ORFs with the GIIM superfamily domain were also present in other orders of Bryopsidales, except *O. quekettii,* suggesting that they are widely conserved in Bryopsidales. A portion of the amino acid sequence also showed weak homology with reverse transcriptases of the order Bryopsidales, suggesting that it may function as a reverse transcriptase.

In addition to protein-coding genes, 26 tRNAs and 3 rRNAs were annotated, consistent with a previous report on *B. plumosa* (NC_026795.1). The anticodons of all the 26 tRNA genes were identical (Table S8).

The chloroplast genome of our line was ∼15 kb shorter than the registered genome (NC_026795.1). This was largely because our line had smaller intergenic regions and fewer introns. For example, the intergenic region between *trnG* (*ucc*) and *rrnF* in our line was 1,362 bp, which was much shorter than that of the other line (13,011 bp). NC_026795.1 had an intron and an intronic ORF in the *rrnL* gene, while neither was present in our line.

### Overview of the mitochondrial genome

The mitochondrial genome of our *Bryopsis* sp. line was assembled as a single closed sequence of 356,152 bp, which was much longer than the hitherto-reported longest sequence in green algae (*O. quekettii*: 241,739 bp) (Repetti et al., 2020). There is one report on the mitochondrial genome of *B. plumosa* (Han et al., 2020). However, our sequences were substantially different from registered sequences. Our own survey of the sequences reported by Han et al. strongly suggested that their specimen belong to Ulvales, and not Bryopsis (Fig. S2).

We compared the obtained sequences with those of other green algae (Table S9). The size of the genome (356,152 bp) was much larger than that of any other mitochondrial genome of green algae (second longest was that of *O. quekettii* at 241,739 bp (Repetti et al., 2020)). This was partly attributed to an increase of introns: we identified 72 introns in 17 genes, which was more than in *O. quekettii* (47 introns in 18 genes) or *C. lentillifera* (29 introns in 13 genes). In extreme cases, 17 introns and 18 exons were present in *cox1*, whereas only 11, 5, and 4 introns were found in *cox1* of *O. quekettii*, *C. lentillifera*, and *Ulva* sp., respectively (Melton et al., 2015; Repetti et al., 2020; Zheng et al., 2018). In total, introns occupied 54.1% of the genome, which was higher than that in *O. quekettii* (39.3%) or *C. lentillifera* (43.4%).

Manual annotation revealed 54 protein-coding genes, 17 tRNAs, and 3 rRNAs. The rRNA numbers were similar to those of most other green algae (Table S9). tRNAs corresponding to 15 amino acids were identified, whereas those corresponding to Ala, Cys, Glu, Lys, and Asn were not.

Of the 54 protein-coding genes, seven were not unambiguously assigned as real ORFs because the encoded amino acid sequences did not show homology to proteins with known functions. In contrast, 47 genes encoded proteins that have conserved domains, many of which are required for mitochondrial function, such as NADH:ubiquinone oxidoreductase (complex I; *nad* genes) or ATP synthase (complex V; *atp* genes) (Table S10). The number of *nad* and *atp* genes encoded in the mitochondrial genome varies among green algae; our *Bryopsis* line often had more than the average number. For example, *nad10* and *tatC* have been found in the mitochondrial genome but not in many other green algae species. However, there was also a reverse case: the mitochondrial genome of *O. quekettii*, but not ours, had *atp4* gene (Table S10).

Manual annotation revealed 72 introns in 17 genes. Introns were more prevalent than those in *O. quekettii* (47 introns in 18 genes) or *C. lentillifera* (29 introns in 13 genes) (Table S9). The number of introns was particularly high in *nad5*, *cob*, *cox1* and *atp1*. In extreme cases, 17 introns and 18 exons were identified in *cox1*, whereas only 11, 5, and 4 introns were found in *cox1* of *O. quekettii*, *C. lentillifera*, and *Ulva* sp., respectively (Fig. S7) (Melton et al., 2015; Repetti et al., 2020; Zheng et al., 2018). The mean intron length was 2,676 bp, which was comparable to that of the two Bryopsodales *O. quekettii* (2,022 bp) and *C. lentillifera* (3,126 bp) (Fig. S8). Introns accounted for 54.1% of the mitochondrial genome, which was higher than that in *O. quekettii* (39.3%) and *C. lentillifera* (43.4%) (Table S9).

Interestingly, 14 protein-coding genes were found in the introns of other genes. A tBLASTn search for published green algal mitochondrial proteins (https://ftp.ncbi.nlm.nih.gov/refseq/release/mitochondrion/) identified three ORFs showing homology to the putative LAGLIDADG endonuclease, ten ORFs showing homology to the putative group II intron reverse transcriptase/maturase, and one ORF encoding a putative protein in the introns of *cox1*, *atp1*, and *rnl* (six in *cox1*, five in *atp1*, and three in *rnl*). The introns of *cox1* contain one gene encoding a LAGLIDADG endonuclease and five genes encoding putative group II intron reverse transcriptases/maturases. The encoded LAGLIDADG endonuclease is likely functional because it possesses LAGLIDADG domains at the N– and C-termini that are required for endonuclease activity (Hausner, 2012; Lambowitz and Belfort, 1993). Four ORFs of the putative group II intron reverse transcriptase/maturase contained one or more RT_G2 introns or RT_like superfamily domains, and three of them possessed the Intron_maturas2 superfamily domain, suggesting that these reverse transcriptases are functional (Table S11).

*O. quekettii* also has endonuclease-like protein ORFs and a putative group II intron reverse transcriptase/maturase on the introns of *cox1*, *atp1*, *rns,* and *rnl*. Thus, the mitochondrial genome size of green algae belonging to the order Bryopsidales, including *Bryopsis*, may have increased in accordance with the increased number and size of introns compared with the mitochondrial genomes of other green algae.

The alignment of *nad2*, *nad7*, *nad5*, *nad9* genes with several green algae, including *O. quekettii* and land plants (*A. thaliana* and *P. patens*), suggested that UGA encodes Trp rather than a termination codon (Fig. S9). This is consistent with other green algae, including *O. queketti*, *Pedinomonas minor*, and *Pycnococcus provasolii* (Noutahi et al., 2019; Repetti et al., 2020).

## Figure legends

**Figure S1.**
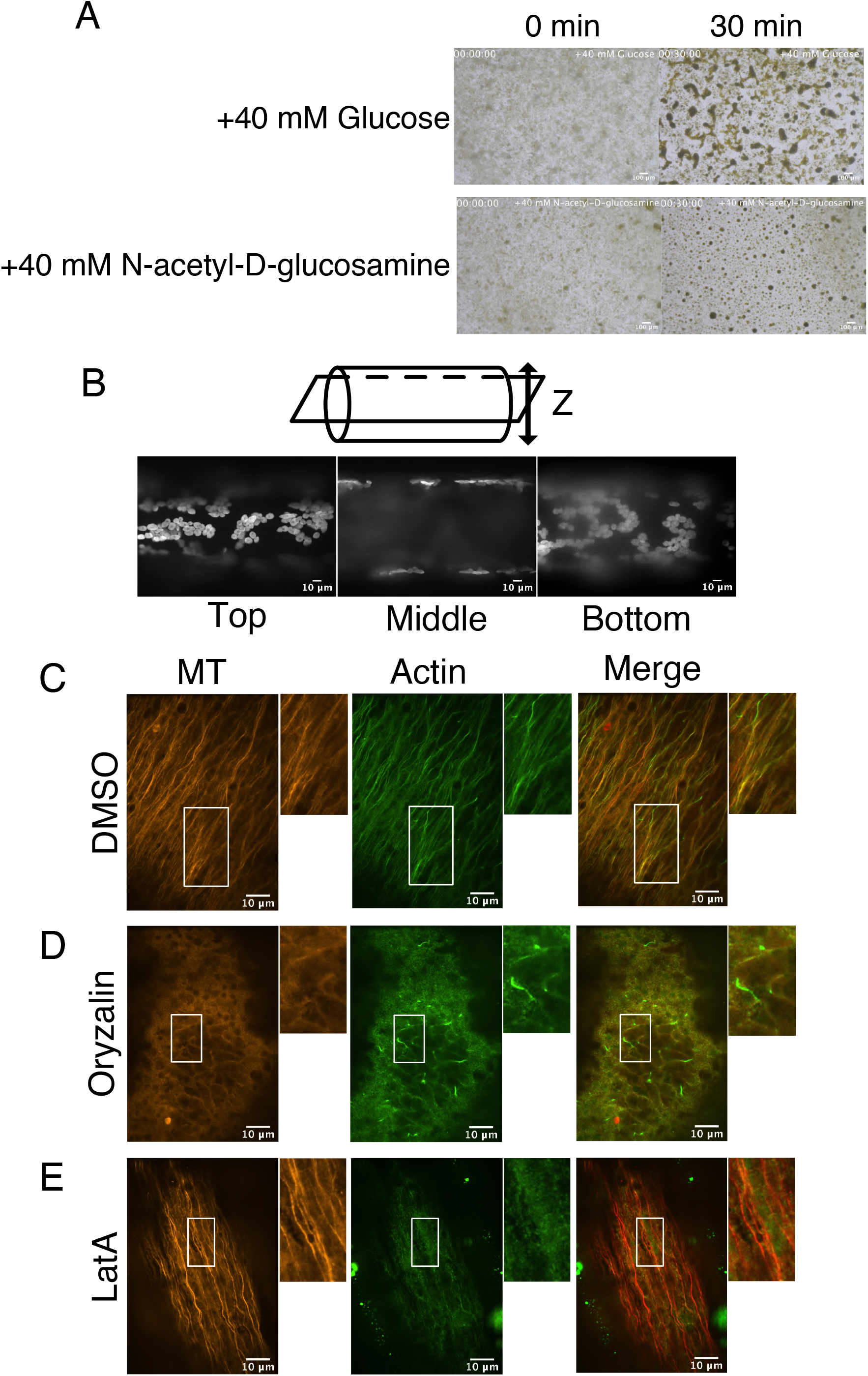
Microtubule and actin organisation in the cytoplasm. (A) Suppression of aggregation of the cytoplasmic extract by N-acetyl-D-glucosamine. Glucose was used as the control. (B) (Top) Schematic representation of the focal plane in microscopy. (Bottom) Three images acquired with 637 nm laser, each representing top, middle, or bottom section of the main axis. Autofluorescent chloroplasts are visualised. A large vacuole occupies the majority of the middle section. (C–E) Immmunostaining of microtubules and actin filaments in the main axis of thalli in the presence or absence of oryzalin (10 µM) or latrunculin A (LatA, 10 µM). The control sample was treated with DMSO. Boxed regions are magnified on the right.

**Figure S2.**
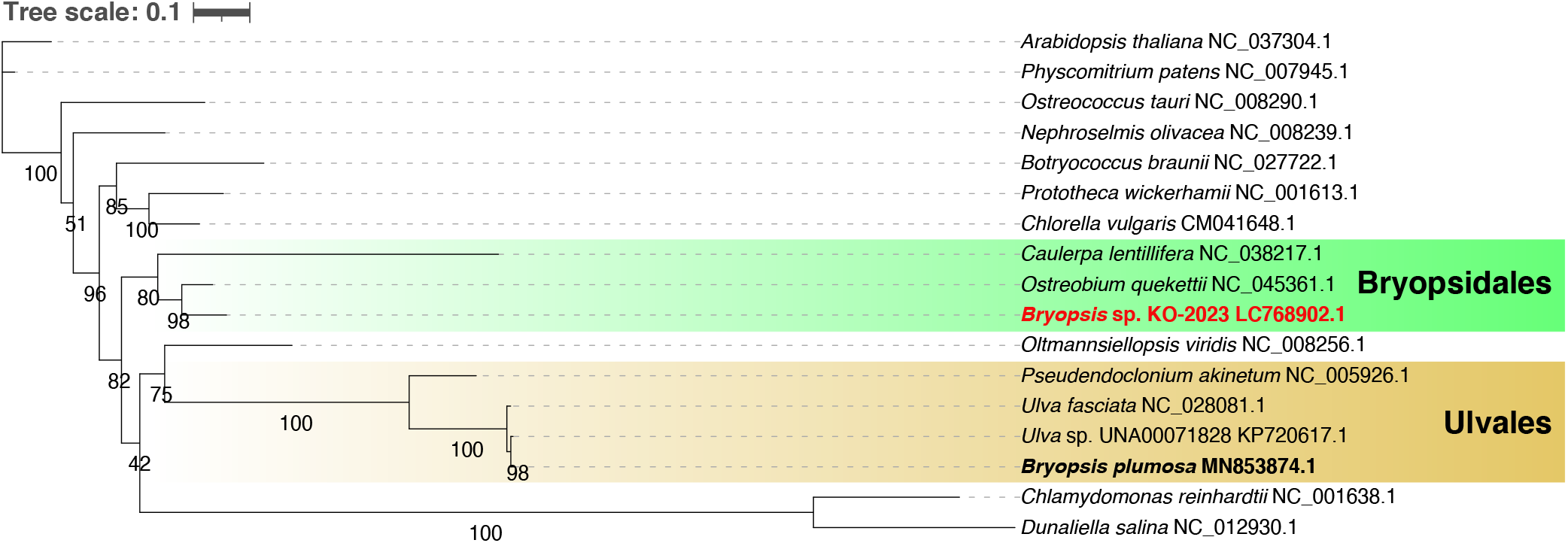
Phylogenetic tree based on mitochondrial genes. *Bryopsis* sp. formed a clade with other Bryopsidales species, whereas the registered ‘*Bryopsis pulmosa*’ sequences (MN853874.1) were most similar to Ulvales sequences. ML gene tree was drawn using IQ-TREE v1.6.12 with LG+F+R4 selected as the best-fit model and branch support was estimated with 1,000 ultrafast bootstrap. The bar indicates 0.1 amino acid substitutions per site.

**Figure S3.**
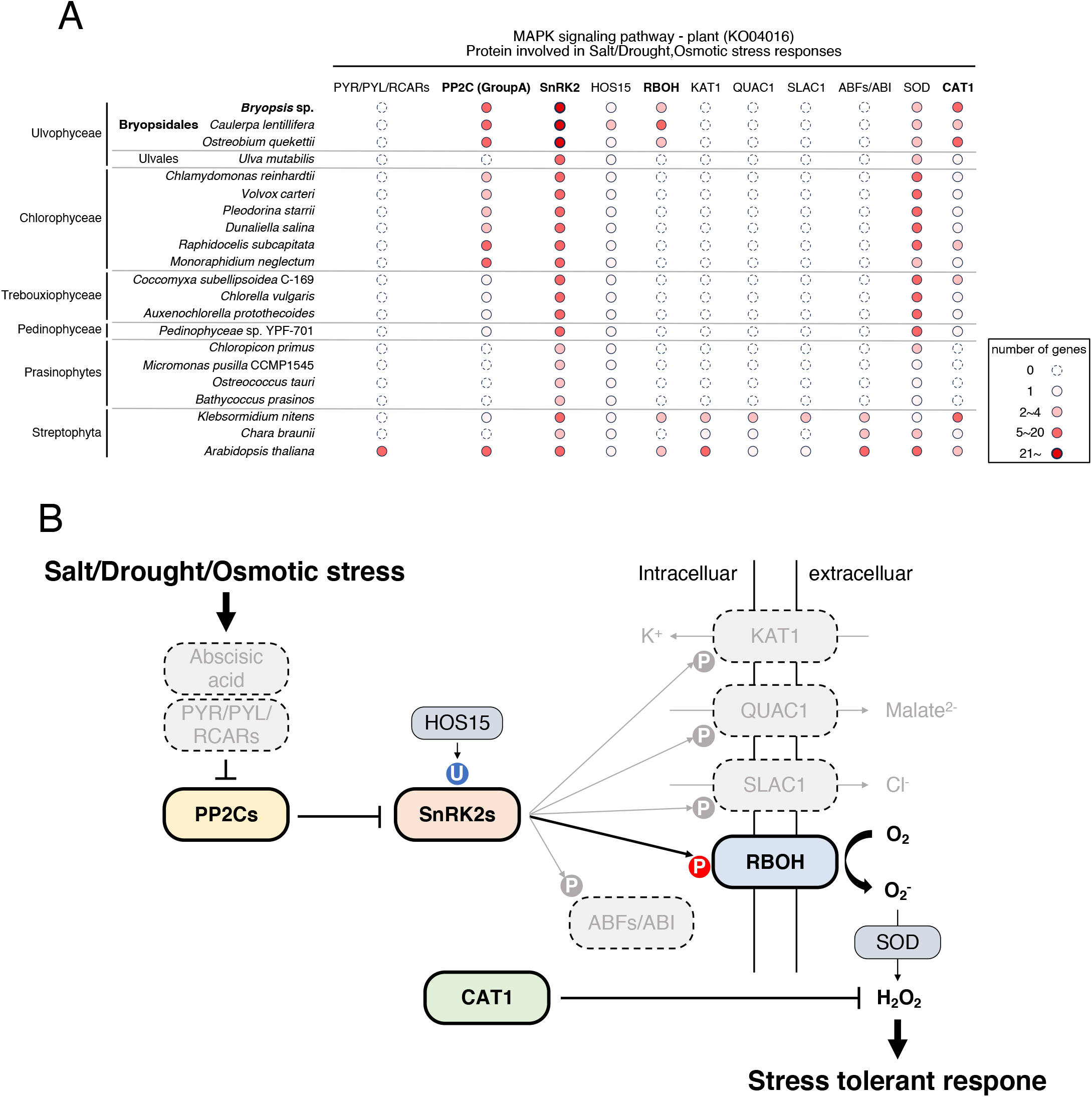
Overrepresenting gene pathway in Bryopsidales. (A) Number of the genes in ‘MAPK signaling pathway – plant (KO04016)’. (B) Signal transduction pathway known in land plants. Figures are derived from ‘MAPK signaling pathway – plant (KO04016)’ in KEGG.

**Figure S4.**
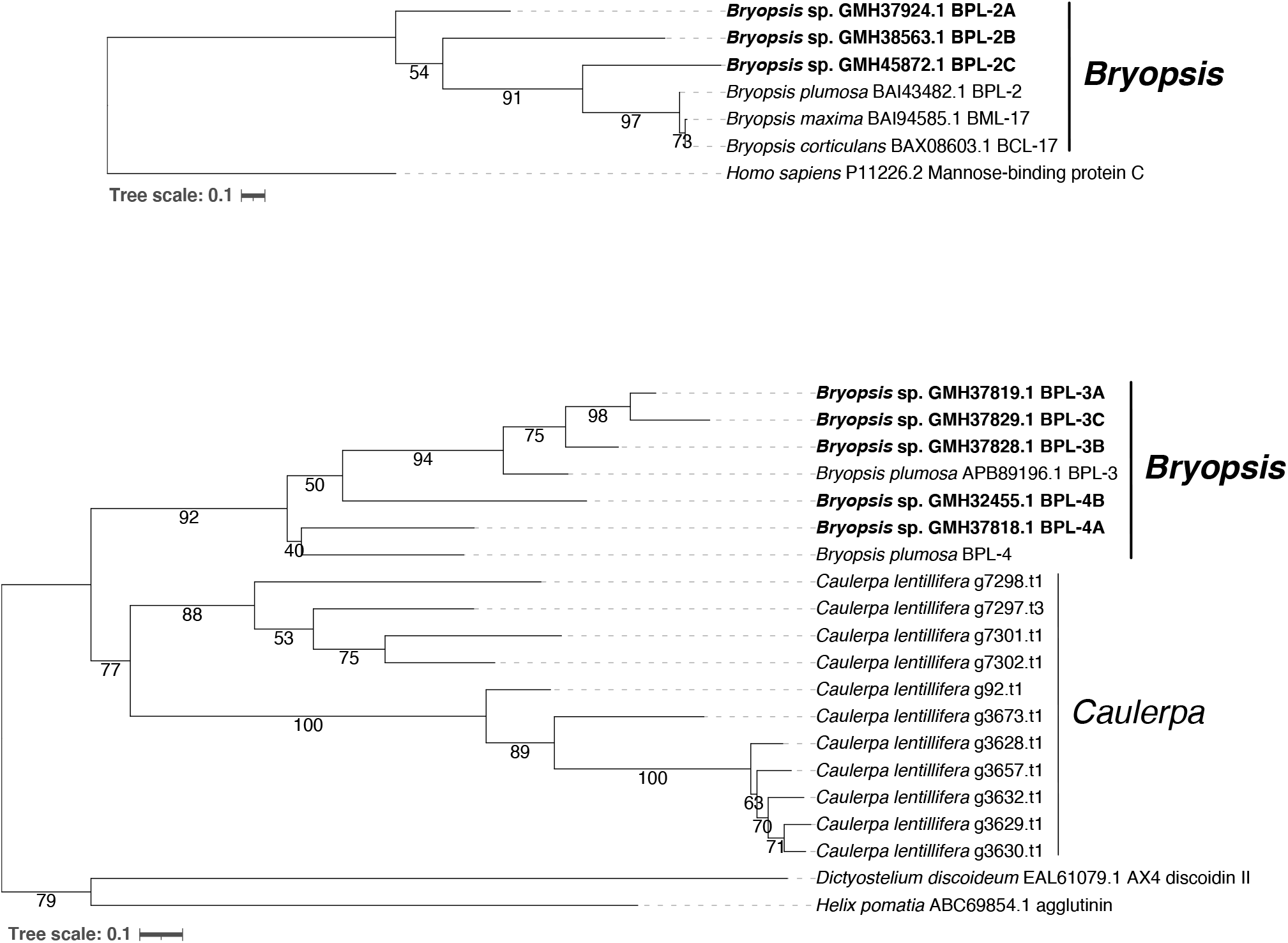
Phylogenetic tree of *BPL-2, 3, 4* genes. ML tree was drawn using IQ-TREE v1.6.12 with WAG+G4 (BPL-2) or LG+G4 (BPL-3/4) selected as the best-fit model and branch support was estimated with 1,000 ultrafast bootstrap. The bar indicates 0.1 amino acid substitutions per site.

**Figure S5.**
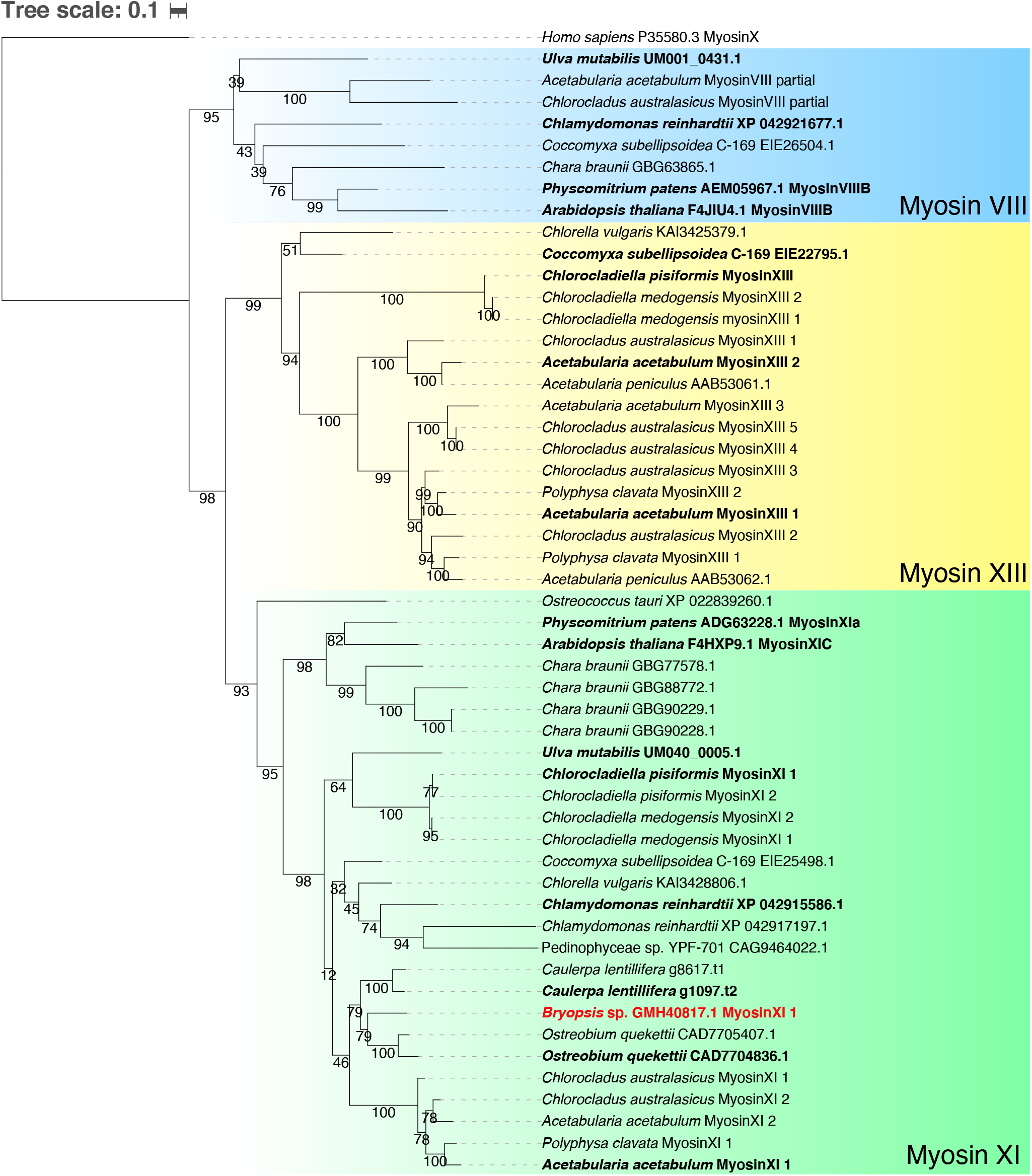
Phylogenetic tree of myosin of green algae. ML tree was also drawn using IQ-TREE v1.6.12 with LG+I+G4 selected as the best-fit model and branch support was estimated with 1,000 ultrafast bootstrap. The bar indicates 0.1 amino acid substitutions per site.

**Figure S6.1.**
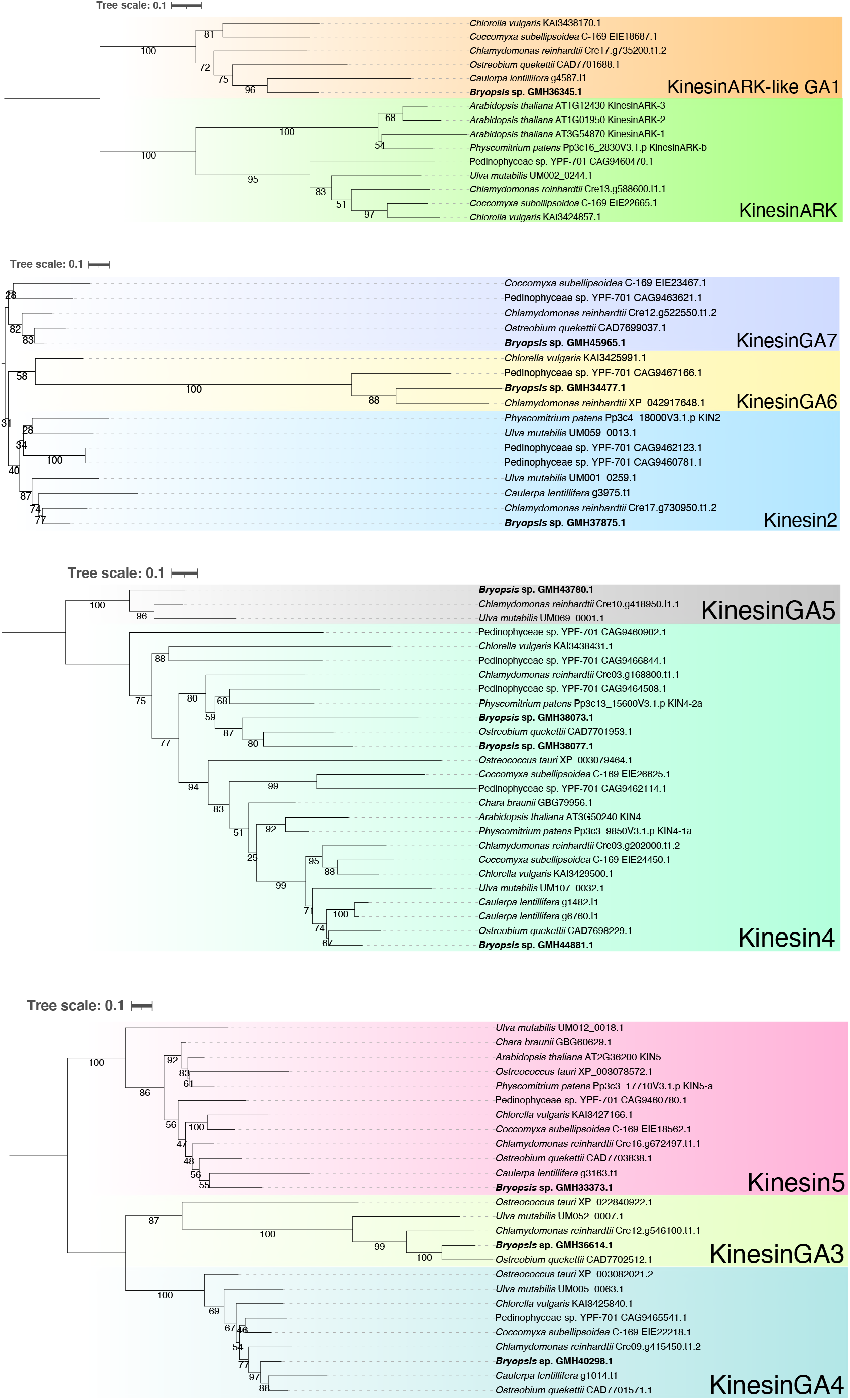
Phylogenetic tree of the kinesin superfamily of green algae. Each page contains trees of a few kinesin subfamilies. Kinesin-GA is alga-specific subfamily. ML tree was also drawn using IQ-TREE v1.6.12 with LG+I+G4 and branch support was estimated with 1,000 ultrafast bootstrap. The bar indicates 0.1 amino acid substitutions per site.

**Figure S6.2.**
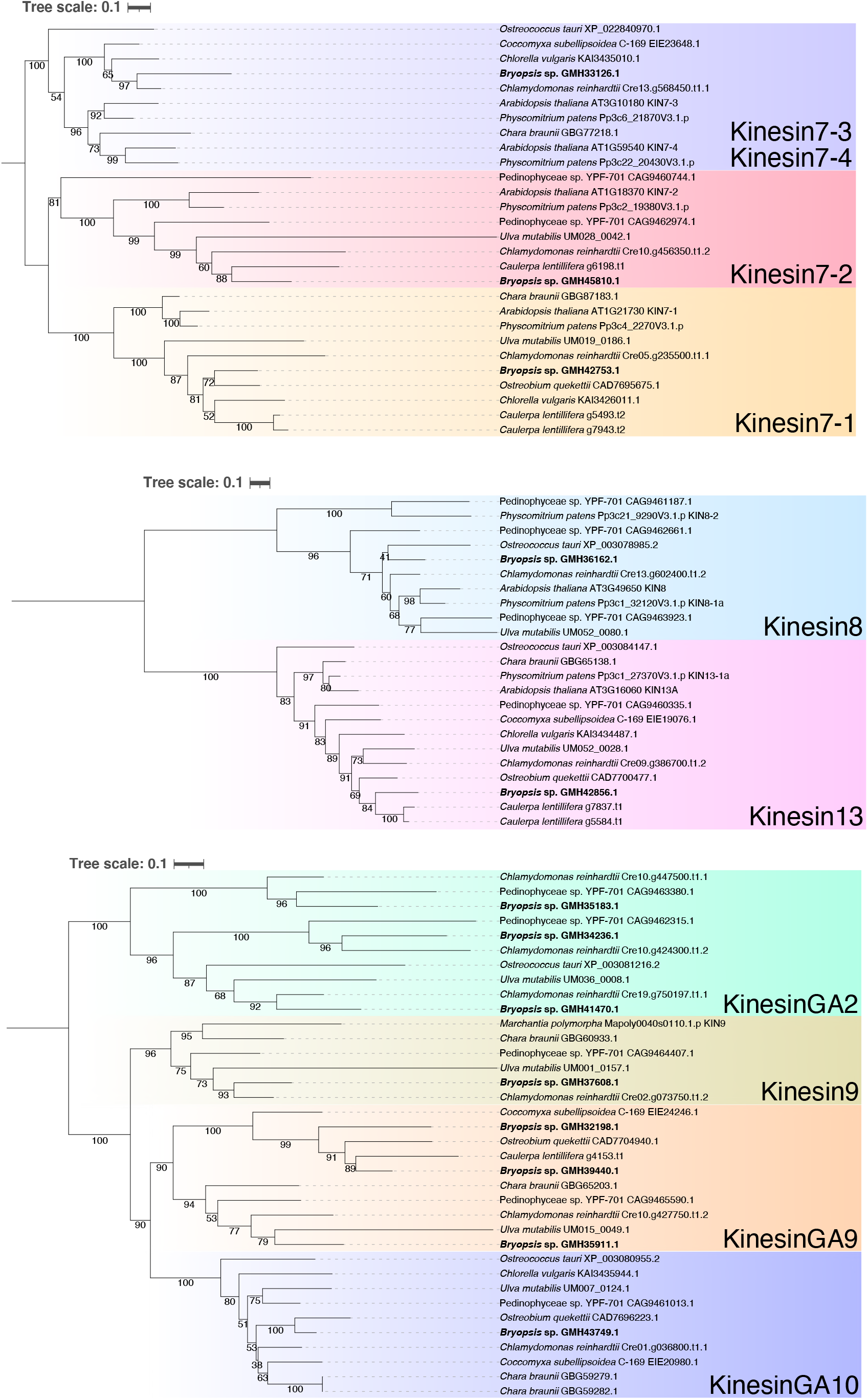
Phylogenetic tree of the kinesin superfamily of green algae. Each page contains trees of a few kinesin subfamilies. Kinesin-GA is alga-specific subfamily. ML tree was also drawn using IQ-TREE v1.6.12 with LG+I+G4 and branch support was estimated with 1,000 ultrafast bootstrap. The bar indicates 0.1 amino acid substitutions per site.

**Figure S6.3.**
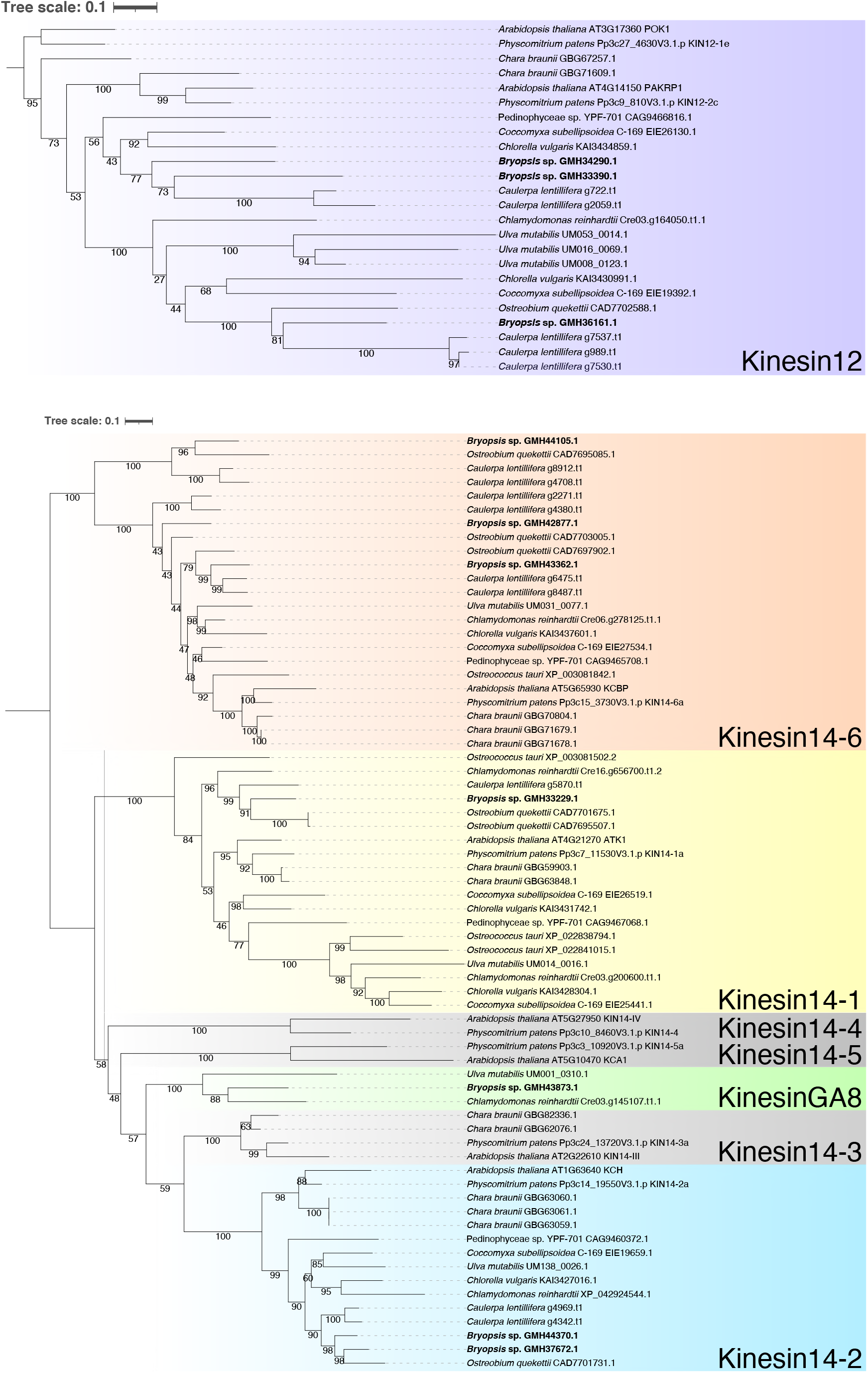
Phylogenetic tree of the kinesin superfamily of green algae. Each page contains trees of a few kinesin subfamilies. Kinesin-GA is alga-specific subfamily. ML tree was also drawn using IQ-TREE v1.6.12 with LG+I+G4 and branch support was estimated with 1,000 ultrafast bootstrap. The bar indicates 0.1 amino acid substitutions per site.

**Figure S7.**
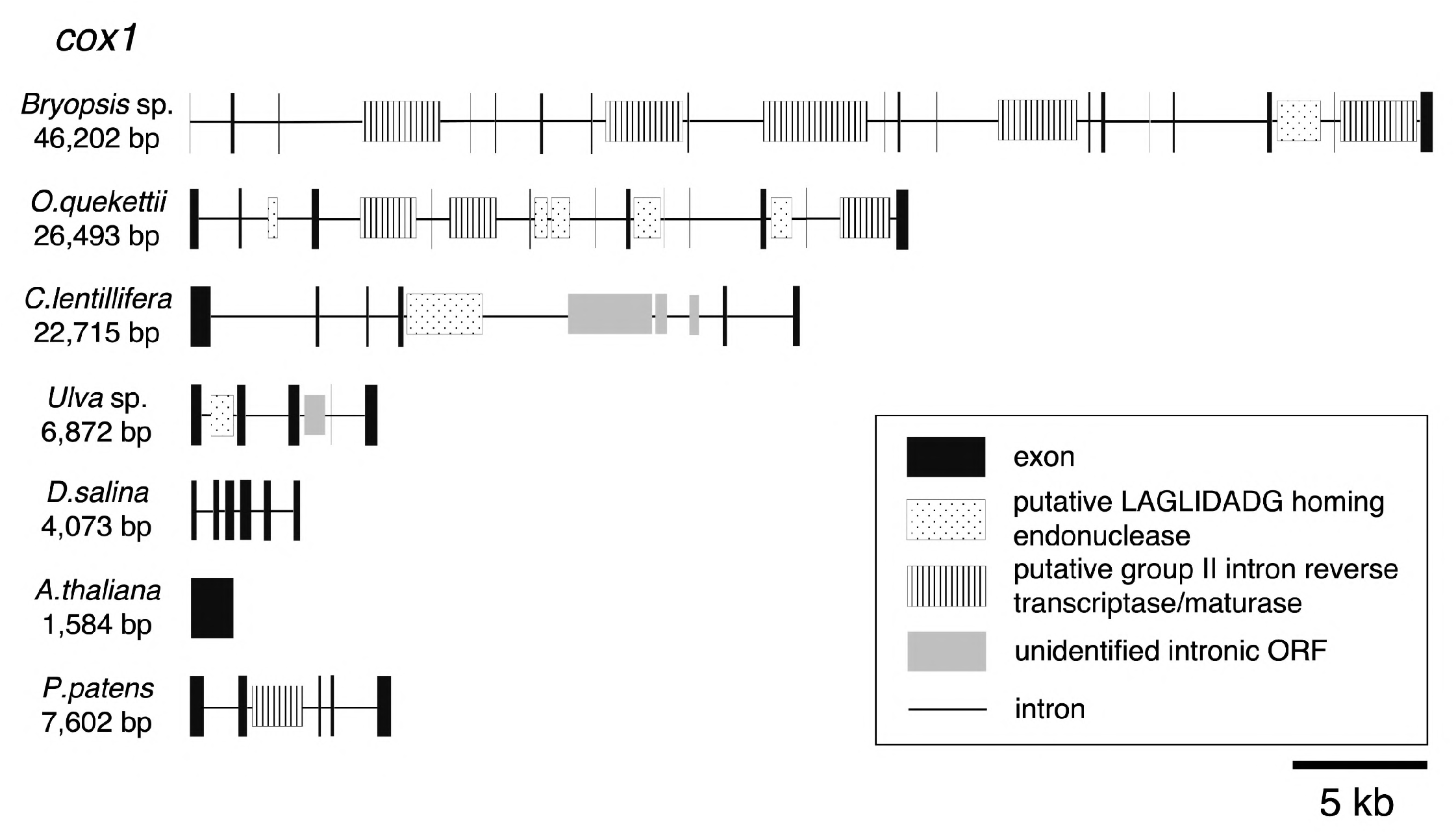
Structure of *cox1* gene encoded in the mitochondrial genome. Several ORFs were identified in the intron of *cox1* gene in *Bryopsis* sp.

**Figure S8.**
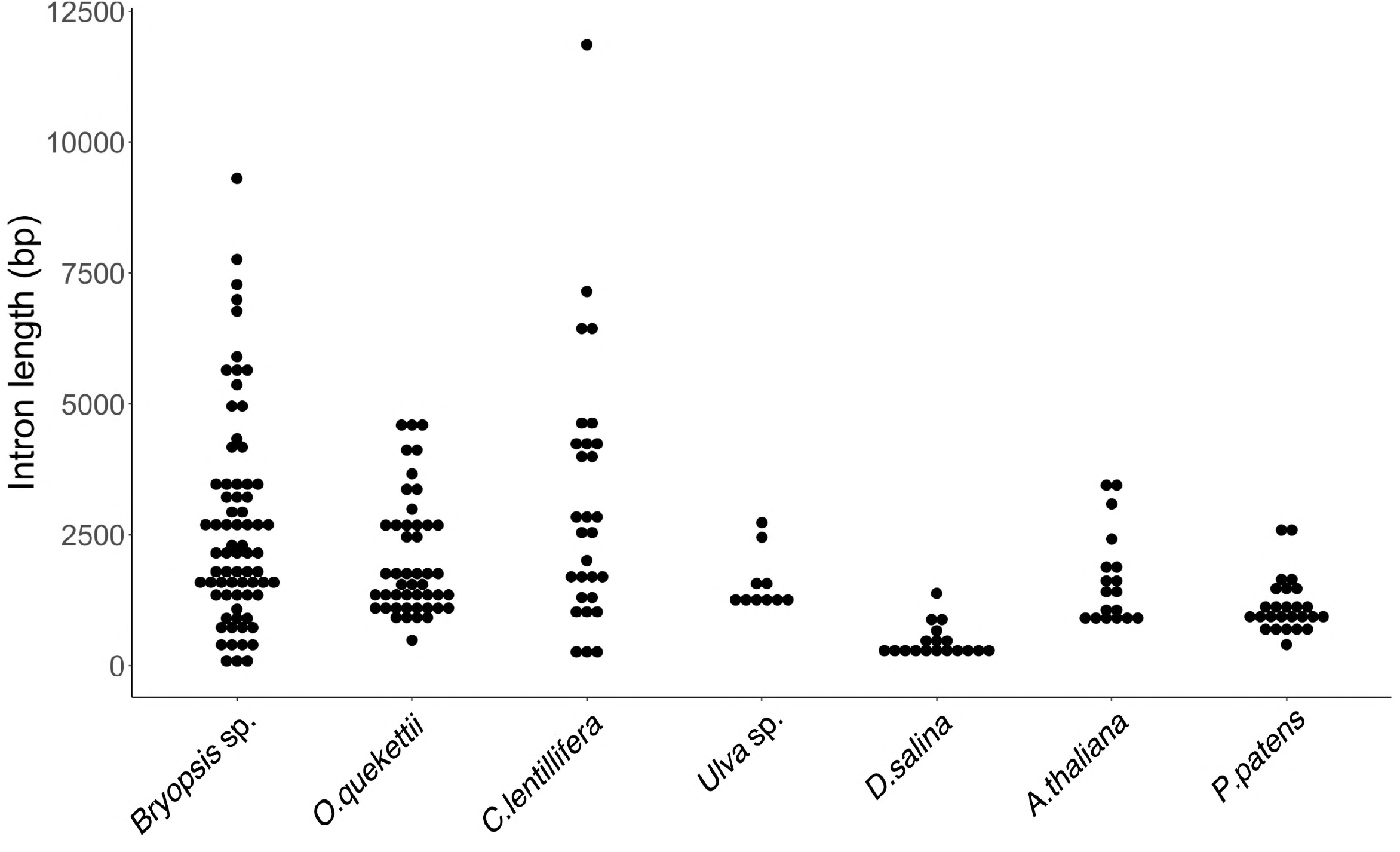
Length of intron in the mitochondrial genome. N = 72, 47, 29, 10, 18, 18, 26 (from left to right).

**Figure S9.**
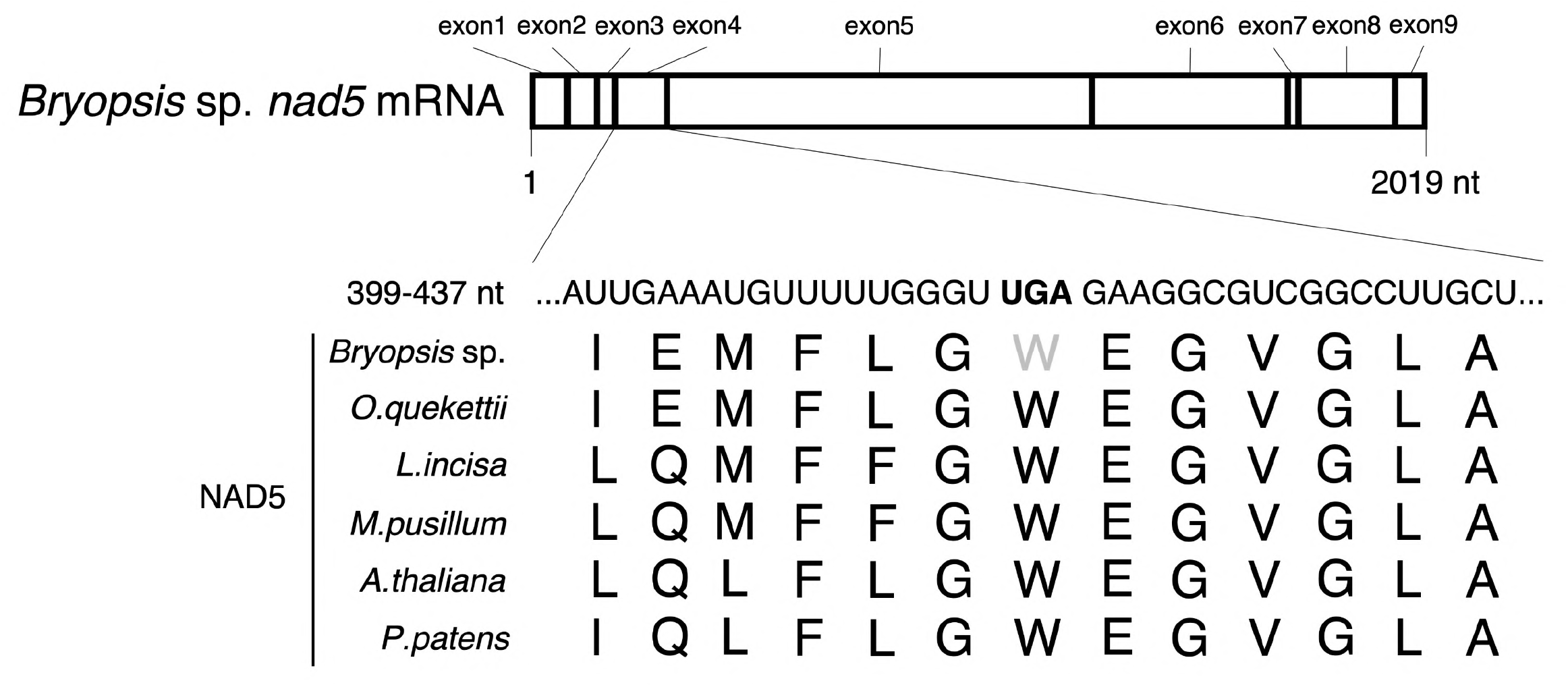
UGA codon likely encodes tryptophan in the mitochondrial genome. Based on the amino acid sequences of the Nad5 protein (this figure) and other conserved proteins in green algae, the UGA of *Bryopsis* sp. likely represents a tryptophan codon, not a termination codon, in the mitochondrial genome.

## Movie legends

**Movie 1. Protoplast formation from extruded cytoplasm**

Images were acquired using a stereomicroscope every 20 s immediately after the extrusion of the cytoplasm into seawater.

**Movie 2. Chloroplast motility in the presence or absence of oryzalin or latrunculin A** Images were acquired every 10 s using a spinning-disc confocal microscope and a 40× 0.95 NA objective lens. Drugs or control DMSO were added at 2 min.

## Supplementary tables

**Table S1. Comparison of the genomes of green algae and land plant species**.

**Table S2. Genome and transcriptome data used in the comparative analysis.**

**Table S3. Number of unigenes based on KEGG pathway annotation.**

**Table S4. Number of genes in each species.**

**Table S5. BUSCO values after transcriptome assembly for Dasycladales and Cladophorales.**

**Table S6. Transcriptome results in the side branch, main axis and rhizoid.**

**Table S7. Comparison of the chloroplast genome of Chloroplastida including *Bryopsis***.

**Table S8. Comparison of protein coding and ribosomal RNA genes encoded in the chloroplast genomes of Chloroplastida including Bryopsis**.

**Table S9. Comparison of the mitochondrial genome of Chloroplastida including *Bryopsis***.

**Table S10. Genes encoded in the mitochondrial genome of Chloroplastida including *Bryopsis***.

**Table S11. Protein-coding genes found on the intron of other genes in the mitochondrial genome**.

